# Different gametogenesis states uniquely impact longevity in *Caenorhabditis elegans*

**DOI:** 10.1101/2023.06.13.544885

**Authors:** Amaresh Chaturbedi, Siu Sylvia Lee

## Abstract

Curtailed reproduction affects lifespan and fat metabolism in diverse organisms, suggesting a regulatory axis between these processes. In *Caenorhabditis elegans*, ablation of germline stem cells (GSCs) leads to extended lifespan and increased fat accumulation, suggesting GSCs emit signals that modulate systemic physiology. Previous studies mainly focused on the germline-less *glp-1(e2141)* mutant, however, the hermaphroditic germline of *C. elegans* provides an excellent opportunity to study the impact of different types of germline anomalies on longevity and fat metabolism. In this study, we compared the metabolomic, transcriptomic, and genetic pathway differences in three sterile mutants: germline-less *glp-1*, feminized *fem-3*, and masculinized *mog-3*. We found that although the three sterile mutants all accumulate excess fat and share expression changes in stress response and metabolism genes, the germline-less *glp-1* mutant exhibits the most robust lifespan increase, whereas the feminized *fem-3* mutant only lives longer at specific temperatures, and the masculinized *mog-3* mutant lives drastically shorter. We demonstrated that overlapping but distinct genetic pathways are required for the longevity of the three different sterile mutants. Our data showed that disruptions of different germ cell populations result in unique and complex physiological and longevity consequences, highlighting exciting avenues for future investigations.

## Introduction

Reproduction is an energy-intensive process and can divert resources from somatic maintenance ^1^, and diminished reproduction often correlates with increased lifespan ^2–8^. Studies in *C. elegans* suggest complex regulation involving antagonizing signals from germline and somatic gonad in modulating longevity ^6^. Lipid metabolism is key to resource allocations and appears to couple reproduction and longevity.

In *C. elegans*, the removal of germline stem cells, either by laser ablation or genetic mutation, results in extended longevity and excessive fat accumulation ^6,9^. Extensive studies of the temperature-sensitive germline-less *glp-1(e2141)* mutant reveal drastic changes in fat metabolism, including altered beta-oxidation, fatty acid desaturation, and lipolysis, and implicate the functional involvement of the lysosomal triglyceride lipase LIPL-4/LIPA and the fatty acid oleic acid ^9–13^. Importantly, germline-mediated longevity involves elaborate germline-to-soma interactions. Germline-mediated longevity is dependent on the intestinal actions of several conserved transcription factors, such as DAF-16/FOXO, DAF-12, SKN-1/NRF, NHR-49/PPARa ^6,10,11,13,14^. Despite the tremendous progress, much remains to be learned about how the germline interacts with the intestine to affect longevity.

The development of the *C. elegans* germline has been well studied, and many gene mutations can cause sterility, including those that disrupt mitotic proliferation or meiotic progression in the germline, or those that fail to produce sperms (feminization of germline, FEM), or oocytes (masculinization of the germline, MOG). In this study, we expanded on the earlier studies that largely focused on using germline-less worms to probe the connection between reproduction, fat metabolism, and longevity. We surveyed a variety of sterile mutants and revealed that whereas all types of self-sterile worms accumulated excess fat, they showed different lifespans. Specifically, we revealed that the germline-less mutants live longer as expected, the feminized mutants also live longer at specific temperatures, whereas the masculinized mutants live shorter, suggesting that increased fat storage is not sufficient to extend lifespan. To further understand the molecular basis of the connection between reproduction, fat metabolism and lifespan, we selected three mutants representing three different types of self-sterility as models, and investigated their lipid and transcriptomic profiles, and their genetic interactions with several key factors. We found that the different sterile mutants shared similar lipid profiles and overlapping transcriptomic changes, but also showed significant differences. Interestingly, the transcription factor DAF-16/FOXO emerged as a key mediator of the full longevity of the long-lived germline-less and feminized mutants, but is dispensable for the short-lived masculinized mutant. Our study has uncovered interesting new avenues of investigation to further probe the complex relationship between reproduction, metabolism, and longevity.

## Results

### Different types of self-sterile mutants accumulated excessive lipids but displayed varied lifespans

To explore the effects of germline development and reproduction on fat accumulation and lifespan in *C. elegans*, we investigated 10 candidate genes, that when inactivated result in different types of self-sterility, using both RNAi and mutant analyses (Table 1). Based on WormBase annotation, we tested: *glp-1* and *iff-1*, which regulate germline proliferation; *gld-1* and *pro-1*, which regulate meiosis; *mog-3* and *fbf-1/2*, which regulate oocyte development; *fem-1, fem-3* and *fog-3*, which regulates sperm development. Irrespective of the mode of sterility, RNAi of each of the genes tested induced worms to accumulate higher levels of lipids based on oil red O (ORO) staining (Supplementary Fig. 1a; Table 1). The accumulated lipids in the germline-less *glp-1* mutant have been shown to be key to its extended longevity ^10^. Interestingly, despite the accumulated levels of lipids, at 20 °C, only RNAi of *glp-1* and *iff-1* showed an increase in lifespan, whereas RNAi of *mog-3, fbf-1/2, mpk-1, gld-1* and *pro-1* resulted in shortened lifespan, and RNAi of *fem-1, fem-3* and *fog-3* showed no change in lifespan (Supplementary Fig. 1b; Table 1). We further validated the RNAi findings using the available loss-of-function mutants and found similar lifespan phenotypes (Supplementary Fig. 1c; Table 1).

To further understand the connection between reproduction, longevity, and lipid metabolism, we selected three mutants, representing the different types of self-sterility, including the germline-less *glp-1(e2141)*, the feminized *fem-3(e1996)* and the masculinized *mog-3(q74)* for detailed studies. Since *glp-1(e2141)* is a temperature-sensitive mutant, we raised all mutants at 25°C and monitored their phenotypes. At the elevated temperature, *glp-1(-)* mutants lived longer as expected, *fem-3(-)* mutants also showed longer lifespan, and *mog-3(-)* mutants lived shorter (Fig. 1b, c; Table 2). To further test how different temperatures can affect the lifespans of these strains, we also checked their lifespans at 15°C. We observed similar results where *glp-1(-)* and *fem-3(-)* mutants lived longer and *mog-3(-)* mutants lived shorter, although all strains had a longer lifespan at 15°C compared to at 25°C as expected (Supplementary Fig. 1d). What accounts for the relatively normal lifespan of the *fem-3(-)* mutant at 20°C, but longer lifespan at 25°C or 15°C is currently unknown.

**Fig. 1:**
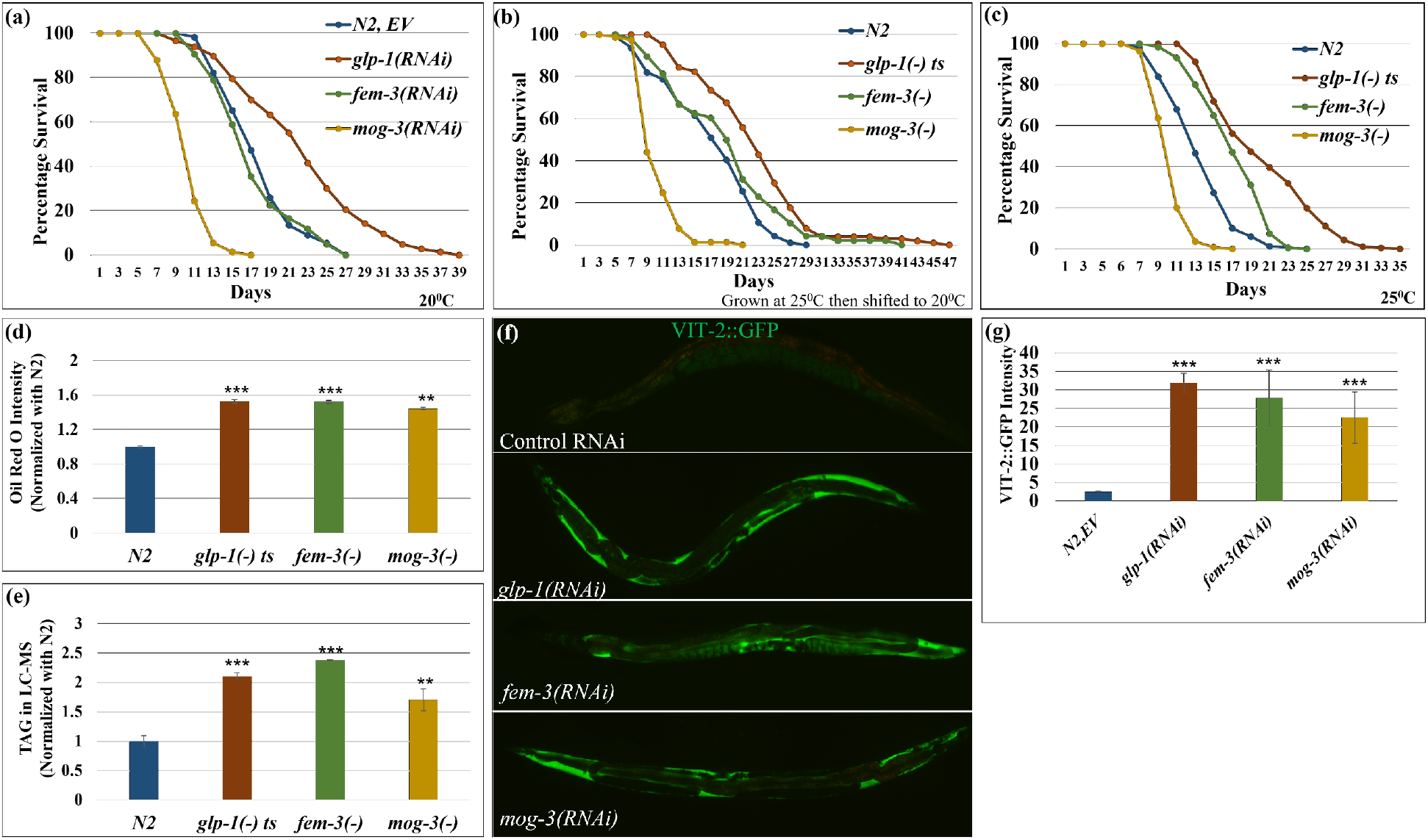
Different modes of sterility induced excessive fat accumulation but differential lifespan. (a). Survival curves showed that *glp-1* RNAi extended lifespans, however, masculinized *mog-3* RNAi shortened lifespan. Feminized *fem-3(RNAi)* worms showed a similar lifespan as wild-type worms at 20°C. (b, c). Survival curves showed that *glp-1(e2141)* and *fem-3(e1996)* worms lived longer, however, *mog-3(q74)* worms lived shorter than wild-type N2 when grown at 25°C then aged at 20°C (b) or grown and aged at 25°C (c). At least 90 worms were scored for each genotype per replicate (N=2). Lifespan data and statistics for a, b and c are shown in Supplementary Table 2. (d-e). Lipid estimated by ORO staining (d) and TAG quantification by LC-MS (normalized with N2) (e) showed that all three sterile mutants *glp-1(e2141), fem-3(e1996)* and *mog-3(q74)* accumulated excess lipid as compared to the wild-type N2 at 25°C. ORO intensity is shown as the mean intensity of two replicates for each genotype with at least 10 worms per replicate. (f, g). Representative images (f) and their quantification (g) showed that excessive VIT-2::GFP accumulated in *glp-1(RNAi), fem-3(RNAi)* and *mog-3(RNAi)* as compared to the wild-type N2 worms. VIT-2::GFP intensity is shown as the mean intensity of two replicates for each genotype with at least 9 worms per replicate. *p< 0.05, **p < 0.01, ***p < 0.001.

We next quantified the lipid levels in the three mutants using different methods at 25°C. As described above, we stained neutral lipids in fixed worms using ORO, imaged the stained worms, and quantified the staining in the first two intestinal cells of each worm (Fig. 1d). We also extracted ORO from stained worms and estimated ORO intensity using colorimetry (Supplementary Fig. 1e). Furthermore, we used a commercial triacylglyceride (TAG) estimation kit to determine the stored TAGs (Supplementary Fig. 1f), which is known to be a preferred storage lipid in worms. Lastly, to survey the lipid composition more broadly in the different sterile mutants, we employed LC-MS (Liquid chromatography-mass spectrometry) to quantify specific lipid species (Fig. 1e). Using these different approaches of lipid estimation, we found that the three sterile mutants accumulated significantly more lipids, particularly TAGs (Fig. 1d, e; Supplementary Fig. 1e, f), at day 1 adulthood. We additionally monitored yolk proteins, which carry lipids from the pseudocoelom to the developing oocytes and represent a major source of nutrients in developing oocytes, using GFP-fused vitellogenin (VIT-2::GFP). We found that all three sterile mutants displayed higher VIT-2::GFP expression as compared to wild-type worms (Fig. 1f, g). In summary, despite their drastically different lifespans, the three sterile mutants accumulated excess lipid and yolk.

### Different types of self-sterile mutants displayed similar but also distinct lipid profiles

We next examined, in more detail, the lipid profiles of the three self-sterile mutants. LC-MS lipidomic analysis detected 1,224 lipid molecules from 15 lipid groups (Fig. 2; Table 3). Principal Component Analysis (PCA) showed that the replicates were highly similar, supporting the high reproducibility of the data. Interestingly, the sterile mutants were clearly separated from wild type (PC1, x-axis, Fig. 2a) and also showed differences among themselves (Fig. 2a). Broadly speaking, the lipidome of each of the tested strains was largely comprised of triacylglycerides (TAG) and phospholipids (PL). Among the phospholipids, the major species were phosphatidylcholine (PC) and phosphatidylethanolamine (PE) (Fig. 2b). Overall, the relative proportion of TAG, PE, and PC remained similar in the three mutants compared to wild-type (Fig. 2b, Supplementary Fig. 2a), with the amount of TAG being significantly upregulated in the sterile mutants, as discussed above (Supplementary Fig. 1f). We further compared the relative levels of the different lipid groups in the different strains and found most of them showed similar levels in the three sterile mutants (Fig. 2c). For instance, phosphatidic acid (PA), glucosylceramide (CerG1), triacylglycerides (TAG), phosphatidylcholine (PCs), and sphingomyelins (SM) were among the upregulated lipid groups in the three sterile mutants compared to wild-type, whereas many other lipid groups did not show substantial change (Fig. 2c). Interestingly, sphingosine (So) levels were lower in *glp-1(-)* and *fem-3(-)* and higher in *mog-3(-)* mutants as compared to wild-type (Fig. 2c).

**Fig. 2:**
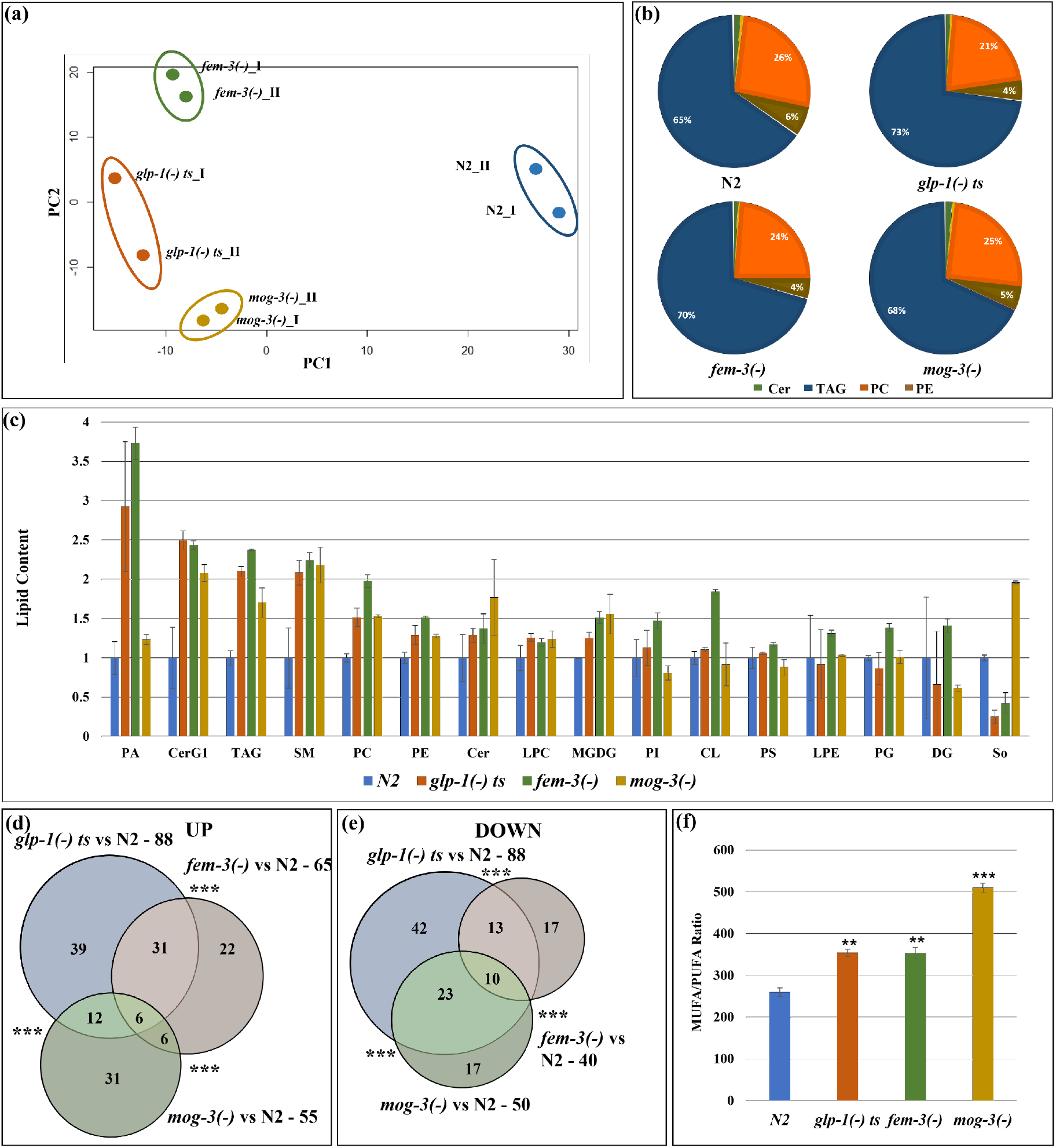
Lipidomic analyses revealed that the different sterile mutant strains share similar but also distinct lipid profiles. (a). PCA analysis of the two replicates of wild-type N2, *glp-1(e2141), fem-3(e1996)* and *mog-3(q74)* showed that the mutants are clearly separated from wild-type N2. Each dot indicates one replicate. PCA plot is generated using EdgeR. (b). Relative distribution of different lipids in N2, *glp-1(e2141), fem-3(e1996)* and *mog-3(q74)* worms showed that the major portion of the lipidome was comprised of TAG, PC and PE and Sphingolipids, especially ceramides. (c). Relative levels of different lipid groups in N2, *glp-1(e2141), fem-3(e1996)* and *mog-3(q74)* worms. The lipid groups were categorized by pooling the normalized counts of the lipid molecules in each of the lipid groups. Y-axis represents the average lipid content of the two replicates in each genotype normalized to wild-type N2. PA – Phosphatidic acid, CerG1 – Glucosylceramide, TAG-Triacylglyceride, SM- Sphingomyeline, PC- Phosphatidylcholine, PE- Phosphatidylethanolamine, Cer- Ceramide, LPC- Lysophosphatidylcholine, MGDG- Monogalactosyldiacylglycerol, PI- Phosphatidylinositol, CL- Cardiolipin, PS- Phosphatidylserine, LPE- Lysophosphatidylethanolamine, PG- Phosphatidylglycerol, DG- Diacylglycerol, So- Sphingosine. (d, e). Venn diagram showing upregulated lipid molecules (d) and downregulated lipid molecules (e) in *glp-1(e2141), fem-3(e1996)* and *mog-3(q74)* vs N2. Venn diagrams were generated using edgeR. A detailed list of lipid molecules is shown in Supplementary Table 3. (f). Ratio of total MUFA (mono-unsaturated fatty acids) and PUFA (poly-unsaturated fatty acids) in wild-type N2, *glp-1(e2141), fem-3(e1996)* and *mog-3(q74)*. The ratios were significantly higher in the sterile mutants compared to wild-type irrespective of their lifespan phenotypes.

We next treated each of the 1,224 detected lipid molecules as a “feature” and used the EdgeR package to identify those that showed statistically significant differences between each of the sterile mutants compared to wild-type worms ^15^. Overall, *glp-1(-)* mutants showed the greatest differences, with 88 upregulated and 88 downregulated lipid species (Fig. 2d, e; Table 3). Comparing the upregulated and downregulated lipid features among the three sterile mutants indicated a significant overlap (Fig. 2d, e; Table 3). We further conducted an enrichment analysis of the significantly changed lipid groups using MetaboAnalyst (www.metaboanalyst.ca) and found that the three sterile mutants shared overrepresentation in some common lipid groups, including TAG as expected. Interestingly, sphingolipids, including sphingomyelins and ceramides, were overrepresented only in *glp-1(-)* and *fem-3(-)* worms (Supplementary Fig. 2c, d, e).

Previous studies have implicated specific fatty acids and elevated MUFA-to-PUFA (monounsaturated fatty acids to polyunsaturated fatty acids) ratio in lifespan modulation ^16–18^. We, therefore, examined the levels of free fatty acids in the three sterile mutants and wild-type worms using LC-MS. We found upregulated MUFA-to-PUFA ratio in all three sterile mutant strains, irrespective of their lifespan phenotypes (Fig. 2f).

In conclusion, we found that the overall lipid compositions were largely similar in the three different sterile mutants at day 1 of adulthood regardless of their lifespan phenotypes. While sterility triggers a common set of overall changes in the lipidome, the sterile mutants also exhibited some differences in lipid profiles, whether those changes contribute to their distinct lifespan phenotypes remains to be investigated.

### Sterile mutants showed significant upregulation of immunity and fat metabolism genes

To understand what could contribute to the lipid and lifespan changes in the sterile mutants at the molecular level, we next compared their gene expression profiles. Previous studies have suggested that the intestine is a key site of action for several major transcription factors that mediate the germline effect on longevity ^10,19,20^. We, therefore, carried out RNA-seq analysis using dissected intestines from the sterile *glp-1(-), fem-3(-)* and *mog-3(-)* mutants and wild-type N2 strain. The intestine-specific transcriptomic analysis also eliminated the difficulty in comparing strains with the variable number of germ cells. We additionally collected whole worms for RNA-seq mainly for comparison with published data ^21^.

Principal Component analysis suggested an overall consistency between biological replicates. The analysis revealed substantial differences in intestinal gene expression between wild-type and sterile mutant strains. Interestingly, the long-lived *glp-1(-)* and *fem-3(-)* strains were separated from wild-type and short-lived *mog-3(-)* strains along PC2 (y-axis, Fig. 3a), suggesting that some gene expression differences could correlate with longevity. We also noted that *glp-1(-)* and *mog-3(-)* worms were separated from wild-type and *fem-3(-)* worms along PC1 (x-axis, Fig. 3a), suggesting that feminized mutants displayed a gene expression profile that is more similar to fully reproductive adults, than to germline-less or masculinized mutants. We also applied PCA to the whole worm transcriptomic data. We found that wild-type and *fem-3(-)* strains grouped closely and were well separated from *glp-1(-)* and *mog-3(-)* strains (PC2, y-axis, Supplementary Fig. 3a), consistent with the observation from the intestinal-specific data. As a quality control, we found that our whole-worm *glp-1(-)* data were highly correlative to the published data (Fig. 3b; Table 4)^21^.

**Fig. 3:**
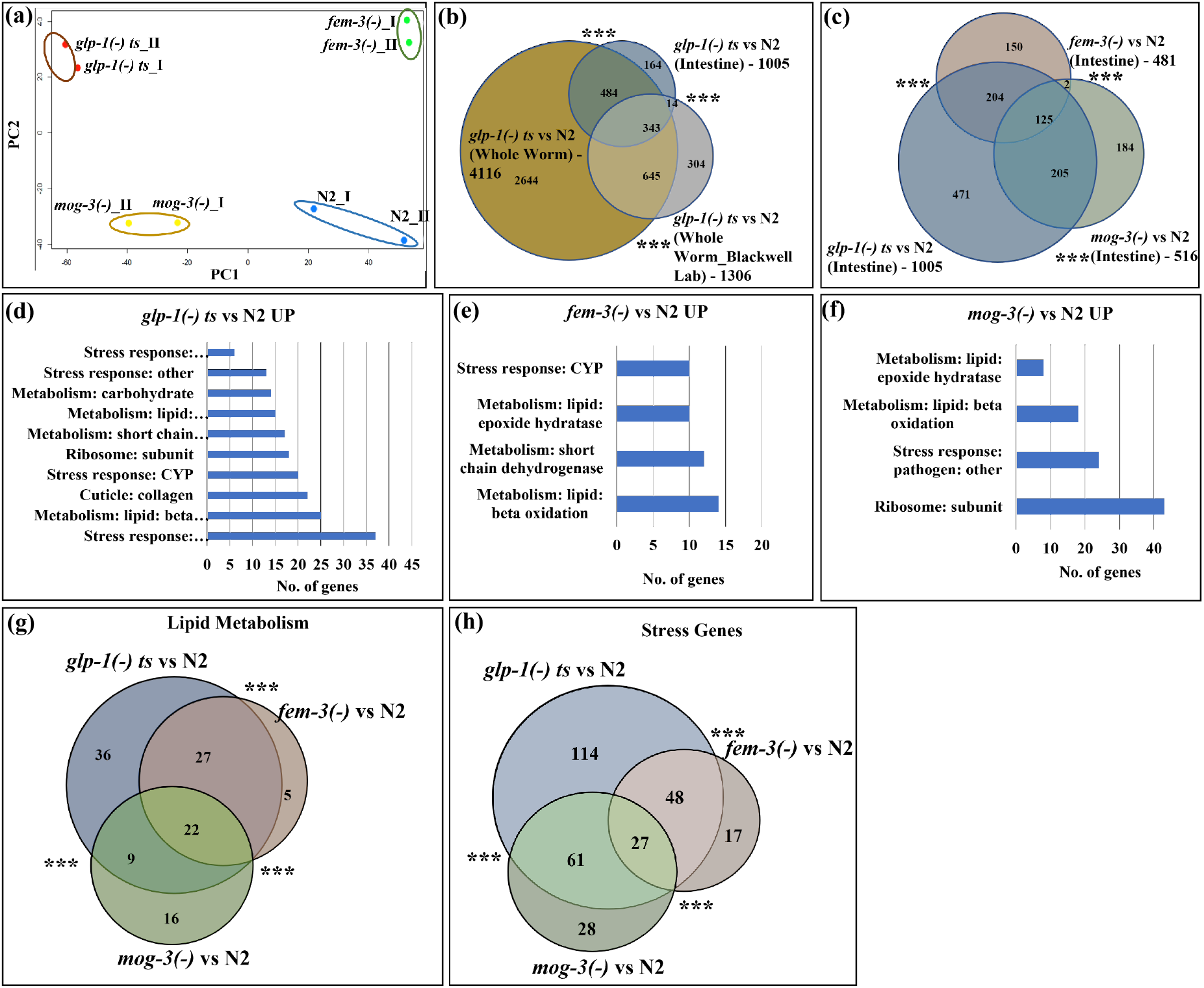
Intestinal-specific transcriptomic analyses revealed that the sterile mutants showed significant upregulation of immunity and fat metabolism genes. (a). PCA analysis of the RNA-seq data for two replicates of N2, *glp-1(e2141), fem-3(e1996)* and *mog-3(q74)* worms showed that the long-lived *glp-1(e2141)* and *fem-3(e1996)* worms are clearly separated from wild-type N2 and short-lived *mog-3(-)* worms. Each dot indicates one replicate. PCA plot is generated using EdgeR. Dissected intestines at the young adult stage were used to generate the RNA-seq data. (b). Venn diagram showed a significant overlap among the upregulated genes in *glp-1(e2141)* worms (relative to N2) in intestine-only, in whole-worm, and in a published data set^21^. The List of genes and overlaps are shown in Supplementary Table 4. (c). Venn diagram showing the significant overlaps between the upregulated genes in *glp-1(e2141), fem-3(e1996)* and *mog-3(q74)*, relative to N2. The *glp-1(-)* mutant showed the greatest gene expression changes (1,005 upregulated genes). Between *fem-3(-)* and *mog-3(-)* mutants, only 125 genes were commonly changed (out of the 481 upregulated genes in *fem-3(-)* and 516 upregulated genes in *mog-3(-*)). The changed genes in *fem-3(-)* and *mog-3(-)* mutants were a subset of the *glp-1(-)* changes (329 of 481 and 330 of 516 in *fem-3(-)* and *mog-3(-)*, respectively). The gene lists and overlaps are shown in Supplementary Table 8. (d, e, f). Gene ontology (GO) terms showed that the metabolism and stress response genes are among the common GO terms that are over-represented in the upregulated genes in *glp-1(e2141)* (d), *fem-3(e1996)* (e), and *mog-3(q74)* (f), relative to N2. Detailed lists of the GO terms are shown in Supplementary Table 9. (g, h). Venn diagram showed the significant overlaps between the upregulated lipid metabolism genes (22) (g)) and stress response genes (27) (h) in *glp-1(e2141), fem-3(e1996)* and *mog-3(q74)*, relative to N2. Venn diagrams were generated using edgeR. ***p < 0.001.

We next identify the genes that showed significant gene expression changes in each of the sterile mutants compared to wild-type. We found 1,005, 481, and 516 upregulated genes and 917, 310, and 456 downregulated genes in the intestinal datasets from *glp-1(-), fem-3(-)*, and *mog-3(-)* mutant strains, respectively (Fig. 3c; Supplementary Table 5). We next compared the intestinal and whole-worm gene expression data for each of the sterile mutants. As expected, the gene expression changes detected in the intestine showed significant overlap with those in whole-worm, and the intestinal changes represented a portion of those detected in whole-worm (Fig. 3c, Supplementary Fig. 3b, c; Supplementary Table 4, Supplementary Table 6 and 7).

To uncover the gene expression differences in the long-lived *glp-1(-)* and *fem-3(-)* and short-lived *mog-3(-)* strains, we compared the significantly upregulated genes in each of the three genotypes. We revealed substantial overlaps among the genes that were upregulated in the three sterile mutants (Fig. 3c; Supplementary Table 8). Interestingly, the germline-less *glp-1(-)* mutant showed the most substantial gene expression changes, whereas the *fem-3(-)* and *mog-3(-)* mutants each exhibited changes that were somewhat distinct and were a subset of the *glp-1(-)* changes (Fig. 3c). Gene ontology (GO) term analysis ^22^ revealed that lipid metabolism and stress response genes were overrepresented among those that showed significant expression changes in all three sterile mutants (Fig. 3d, e, f; Supplementary Table 9-11. Upon comparing the upregulated lipid metabolism and stress genes, 22 fat metabolism genes (Fig. 3g) and 27 stress genes (Fig. 3h) were found to be common among *glp-1(-), fem-3(-)* and *mog-3(-)* mutants. We additionally performed clustering analysis of the significantly changed genes using STRING, a protein network analysis tool ^23^. We again found that the upregulated genes in the three sterile mutants are well-represented for fat metabolism and stress response pathways (Supplementary Fig. 4a, b, c; Supplementary Table 12). Further STRING analysis of the fat metabolism cluster revealed that all three mutants showed expression changes in the sphingolipid metabolism, however, the number of genes in the clusters varies in these mutants; *glp-1(-)* and *fem-3(-)* have 21 and 17 genes, respectively whereas, *mog-3(-)* mutant only have 3 genes (Supplementary Fig. 4d, e, f; Supplementary Table 13). Both GO and STRING analyses also indicated that the sterile mutants showed some unique gene expression changes, including structural and autophagy components in *glp-1(-)* and ribosomal biogenesis in *mog-3(-)* (Supplementary Fig. 4). To gain further insights into the specific fat metabolism pathways that might be altered in the three sterile mutants, we examined the RNA fold change of key enzymes in several major fatty acid metabolism pathways based on our RNA-seq data. This analysis revealed some interesting changes that are generally consistent among the three sterile mutants. Strikingly, the fatty acid desaturase *fat-7* was downregulated in *glp-1(-)* and *fem-3(-)* mutants but upregulated in *mog-3(-)* mutant, and the *acs-2* was greatly downregulated in *mog-3(-)* mutant but not changed in *glp-1(-)* and *fem-3(-)* mutants (Supplementary Fig. 5).

Among the stress response genes that were significantly changed in the three sterile mutants, pathogen/innate immune genes were particularly overrepresented (Fig. 3d, e, f, Supplementary Fig. 4; Supplementary Table 9). We further compared the upregulated genes that were annotated to be pathogen-responsive gene ^22^ among the three sterile mutants and observed a significant overlap, where the changes in *fem-3(-)* and *mog-3(-)* each represented a subset of those changed in *glp-1(-)* mutant (Supplementary Fig. 6a; Supplementary Table 14). These data prompted us to examine the immunity phenotype of the three sterile mutants. We employed the well-established model of *Pseudomonas aeruginosa (PA14)* infection^24^ and found that all three mutants survived better upon PA14 infection compared to wild-type worms (Supplementary Fig. 6b; Supplementary Table 15).

Taken together, the results suggested that sterility induced a shared pattern of gene expression changes and disrupting self-fertilization due to sperm or oocyte loss each produced gene expression changes that represented a subset of those associated with complete loss of the germline.

### Integration of transcriptomic and lipidomic data highlighted the sphingolipid metabolism pathway to show significant alteration in the sterile mutants

Our lipidomic and transcriptomic analyses discussed above revealed interesting changes in the three sterile mutants. We reasoned that connecting the transcriptomic and lipidomic data could provide further biological insights. We next input the genes and lipid molecule that showed significant changes in the sterile mutants into MetaboAnalyst (www.metaboanalyst.ca), a network analysis tool capable of connecting transcriptomic and metabolomic data. A strong signature of the sphingolipid metabolism pathway emerged from this analysis in all three sterile mutants (Fig. 4). Key genes in this pathway showed upregulated expression in all three mutants, with more components showing changes in the *glp-1(-)* mutant (Fig. 4a) compared to the *fem-3(-)* and *mog-3(-)* mutants (Fig. 4b, c). Among the metabolic intermediates, glucosylceramide was commonly downregulated in all three mutants, but sphingosine was downregulated only in *glp-1(-)* and *fem-3(-)* mutants, and ceramide was uniquely upregulated in *fem-3(-)* mutant (Fig. 4). The data highlighted the sphingolipid metabolism pathway to be significantly altered in the three sterile mutants. However, the cause and biological consequence of this alteration need further investigation.

**Fig. 4:**
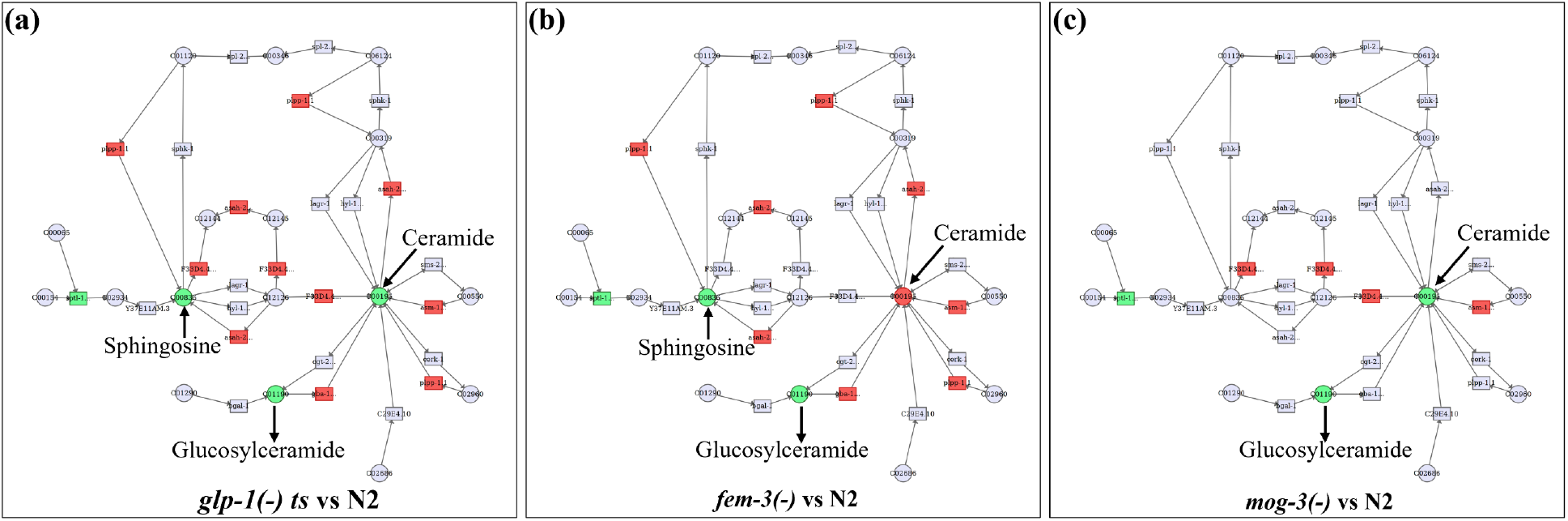
Integration of lipidomic and transcriptomic data. (a-c). The lipid molecules (Fig. 2) and genes (Fig. 3) that showed significant changes in the three sterile mutants compared to the wild-type were input into MetaboAnalyst (www.metaboanalyst.ca). The output data indicated the lipid molecules (circles) and genes (rectangles) that are connected based on published data. Green indicates downregulation and Red indicates upregulation.

### The lifespans of different sterile mutants are dependent on distinct known genetic pathways

Our analyses thus far indicated that *glp-1(-), fem-3(-)* and *mog-3(-)* mutants displayed similarities in altered lipid metabolism and elevated stress response; however, it remains unclear what contributes to their lifespan differences. We next tested whether the three sterile mutants engage different genetic pathways that are known to be key to longevity determination. DAF-16/FOXO is a key downstream transcription factor of insulin/IGF-1 signaling pathway and has been implicated in several longevity-affecting regimens, including lifespan extension mediated by germline loss. Upon germline removal, DAF-16/FOXO enters the intestinal nuclei and regulates downstream target genes ^6^. We tested the effects of *daf-16* removal in wild-type, *glp-1(-), fem-3(-)* and *mog-3(-)* mutant strains. In our experimental condition of 25°C, *daf-16* RNAi completely suppressed the extended lifespan of the *glp-1(-)* and *fem-3(-)* worms but did not significantly alter the already shortened lifespan of the *mog-3(-)* mutant (Fig. 5a; Supplementary Fig. 7a; Supplementary Table 16). We also examined DAF-16::GFP localization in the intestinal cells of the various strains, which has previously been shown to become nuclear localized in germline-less worms ^19^. We found that most of the *glp-1(-)* worms showed nuclear DAF-16::GFP, as did *fem-3(-)* worms; in contrast, DAF-16::GFP remained cytoplasmic in *mog-3(-)* worms (Fig. 5d, e). We additionally confirmed these results using SOD-3::GFP, a reporter of DAF-16 activation ^25^. As expected, *glp-1(-)* worms displayed increased SOD-3::GFP intensity ^26^. Interestingly, we found that *fem-3(-)* worms also showed high induction of SOD-3::GFP, whereas *mog-3(-)* worms did not display any change in SOD-3::GFP intensity (Supplementary Fig. 7b, c). We additionally performed a time-course experiment to assess the timing of DAF-16 activation. We found that DAF-16::GFP localized to intestinal nuclei in *glp-1(-)* and *fem-3(-)* worms starting in day 1 adult (1 DA) and persisted to day 12 (12 DA) (Fig. 6a). In contrast, *mog-3(-)* worms showed slight nuclear localization of DAF-16::GFP in day 1 adult worms but not at the other time points examined (Fig. 6a). Interestingly, we found that DAF-16 became nuclear localized starting in day 5 adulthood in wild-type worms and remained nuclear at the last time point (Fig. 6a). These data corroborated with earlier data (Fig. 5, Supplementary Fig. 7) and suggested that the persistent activation of DAF-16 starting on day 1 adulthood could be essential to lifespan extension.

**Fig. 5:**
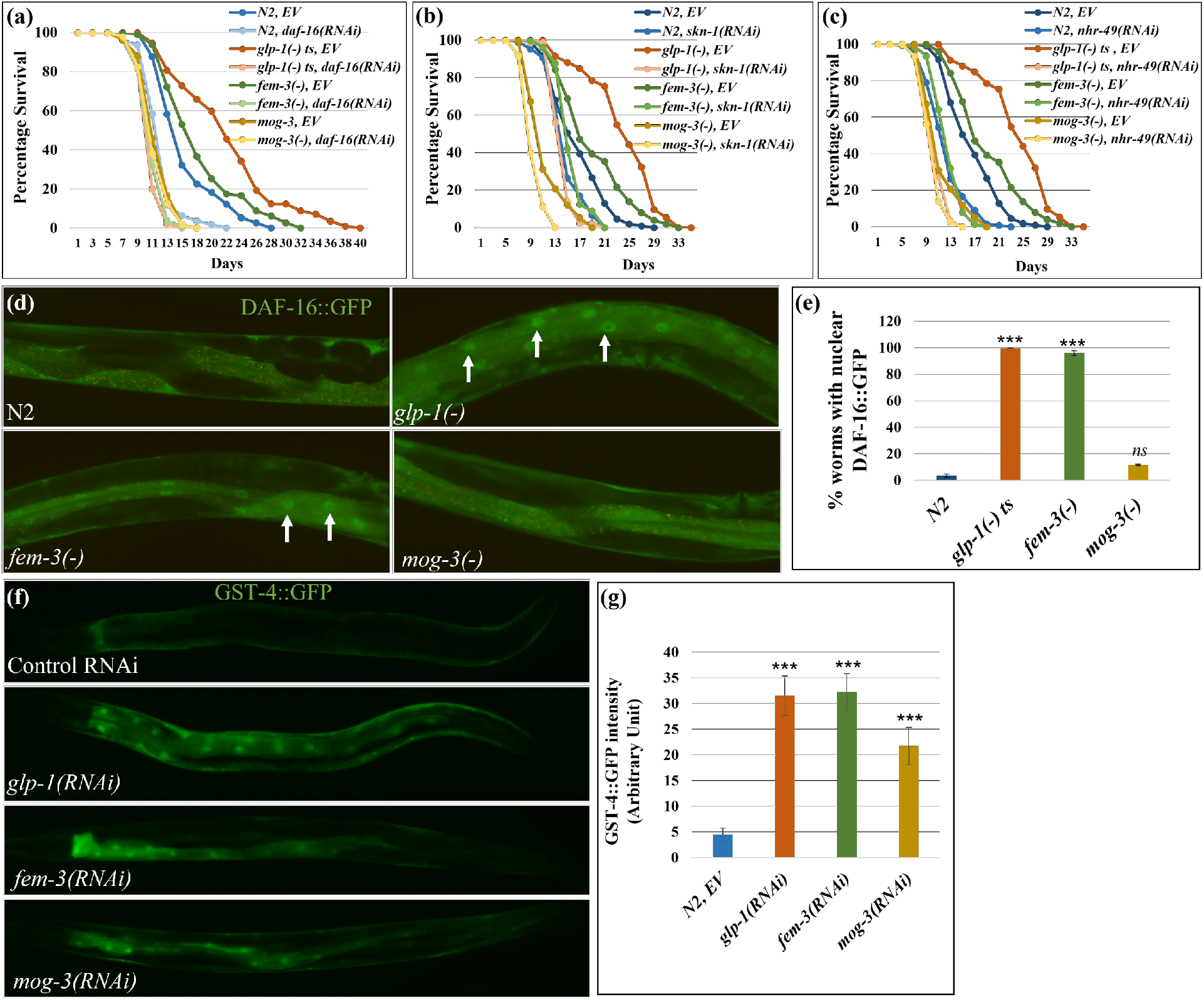
The lifespans of different sterile mutants were dependent on distinct genetic pathways. (a-c) Survival curves showing the lifespans of wild-type N2, *glp-1(e2141), fem-3(e1996)*, and *mog-3(q74)* worms upon *daf-16* RNAi (a), *nhr-49* RNAi (b), and *skn-1* RNAi (c) (lifespan data and statistics are shown in Supplementary Table 16). Lifespan assays were done using at least 90 worms for each genotype per replicate (N=2). (d, e). Representative images (d) and quantification (e) showed that nuclear localization of *daf-16::gfp* is elevated in *glp-1(RNAi)* and *fem-3(RNAi)* worms but not in *mog-3(RNAi)* worms. Y-axis represents mean GFP intensity of two replicates of each genotype with at least 10 worms per replicate. ***p < 0.001, ns - not significant. Arrowheads mark nuclear DAF-16::GFP. (f, g). Representative images (f) and their quantification (g) showed that *gst-4::gfp* is elevated in *glp-1(RNAi), fem-3(RNAi)* and *mog-3(RNAi)* worms. Y-axis represents mean GFP intensity of two replicates for each genotype with at least 10 worm per replicate. ***p < 0.001.

**Fig. 6:**
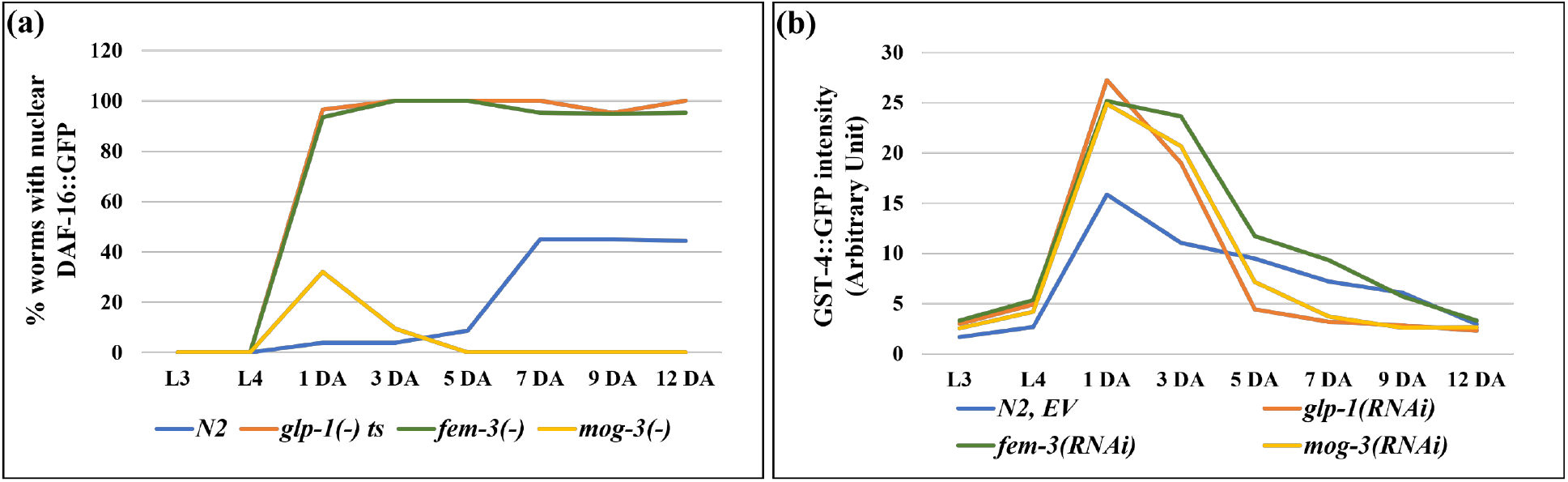
Time course analysis of DAF-16 and SKN-1 activation. (a). Percent wild-type N2, *glp-1(e2141), fem-3(e1996)* and *mog-3(q74)* worms with nuclear DAF-16::GFP at 8 different stages of their lives, including two larval stages (L3 and L4), two reproductive stages (day 1 and 3 of adulthood, 1 DA and 3 DA), and four post-reproductive stages (5 DA, 7 DA, 9 DA, 12 DA). (b) GST-4::GFP signal intensity in wild-type N2 treated with the indicated RNAi at 8 different stages of their lives as described previously. At each time point, two replicates of each genotype were scored with at least 50 worms per replicate. These experiments were performed at 25°C.

Excess fat and yolk have been suggested to activate the transcription factor SKN-1/Nrf2 and promote lifespan in germline-less mutants ^21^. Additionally, SKN-1 has been shown to be activated in several long-lived mutants ^21,27,28^. Since the three sterile mutants accumulated elevated levels of lipid and vitellogenin (Fig. 1, Supplementary Fig. 1), we next tested whether their lifespans require functional SKN-1. We found that the lifespans of all four strains (wild type, *glp-1(-), fem-3(-)* and *mog-3(-)*) were decreased upon depletion of *skn-1* (Fig. 5b, Supplementary Fig. 7a). The degree of lifespan reduction was greatest in the *glp-1(-)* mutant, and least in the short-lived *mog-3(-)* mutant (Supplementary Fig. 7a; Supplementary Table 16). We additionally used GST-4::GFP as a reporter of SKN-1 activity ^29^. We found that *glp-1(-)* worms displayed increased GST-4::GFP intensity as expected; interestingly, *fem-3(-)* and *mog-3(-)* worms also showed increased expression of GST-4::GFP (Fig. 5f, g). We additionally used GST-4::GFP to monitor SKN-1 activation over time in the three sterile mutants. We observed the onset of GST-4::GFP induction at the L4 stage, which persisted until day 7 adulthood, with peak elevation in 1-day-old adults in all the strains tested (Fig. 6b). GST-4::GFP activation was greater in the sterile mutants compared to wild-type, consistent with these mutants having elevated levels of lipid. Overall, these results indicated that SKN-1 is activated in all three sterile mutants, and appears essential for their full longevity, despite that they have very different lifespans.

Nuclear hormone receptor (NHR) signaling pathways have also been shown to modulate longevity ^10,13,30^. NHR-49/PPARα in particular is a key transcription factor that regulates fatty acid metabolism and affects lifespan; *nhr-49* depletion in *C. elegans* causes shortened lifespan and elevated fat content ^30^ and also suppresses the longer lifespan of *glp-1(-)*^13^. We next tested the effect of *nhr-49* loss in the sterile mutants and found that the lifespans of all the strains tested, including wild-type, were decreased by the removal of *nhr-49* (Fig. 5c, Supplementary Fig. 7a). *nhr-49* RNAi had the greatest lifespan shortening effect on *glp-1(-)* mutant and completely suppressed the lifespan increase of the *fem-3(-)* mutant (Supplementary Fig. 7a; Supplementary Table 16). These results indicated that NHR-49 is likely essential for the full longevity of the sterile mutants, despite their different lifespans.

### DAF-16 and SKN-1 activation in long-lived *glp-1(-)* and *fem-3(-)* are differentially dependent on accumulated fat

Previous studies have suggested that altered fat levels could induce longevity-promoting factors ^9,10^. Therefore, we investigated the roles of fat accumulation on DAF-16 and SKN-1 activation in the three sterile mutants. FAT-6 and FAT-7 encode fatty acid desaturases and are key to the production of MUFAs and overall fat storage. Since *fat-6* and *fat-7* share significant sequence homology, they can be knocked down by a single RNAi. We found that nuclear accumulation of DAF-16::GFP was partially but significantly rescued in *glp-1(-)* and *fem-3(-)* worms upon RNAi depletion of *fat-6/fat-7*. Nuclear DAF-16::GFP was detected in ∼40% and ∼23% of the worms in *fat-6* RNAi-treated *glp-1(-)* and *fem-3(-)* worms, respectively, compared to ∼90% in control RNAi-treated worms (Supplementary Fig. 8a and 9).

Interestingly, fat depletion completely reversed the intestinal expression of GST-4::GFP in all sterile worms, including the faint expression in WT worms (Supplementary Fig. 8b and 10). The results indicated that excess fat storage is key to intestinal SKN-1 activation in the strains tested, irrespective of their lifespan phenotype. Furthermore, DAF-16 activation partially depended on fat storage, but additional factors likely also contribute to DAF-16 activation in *glp-1(-)* and *fem-3(-)* mutants.

## Discussion

In this study, we investigated different types of self-sterility in *C. elegans* to better understand the connection between reproduction, fat metabolism, and longevity. Our results revealed that different types of sterility result in the accumulation of excess lipids, but only defective proliferation of germline stem cells leads to robust lifespan extension, similar to previous studies ^6^. We further showed that the different sterile mutants share similar overall lipid profiles but also some significant differences (Fig. 2 and Supplementary Fig. 2). Due to the need of a substantial amount of starting material, we carried out LC-MS lipid profiling using whole worms. Although the three sterile mutants accumulate higher levels of triglycerides (∼2-fold higher than wild-type), the relative proportion of triglycerides to phospholipids remains similar between the *fem-3(-), mog-3(-)*, and wild-type worms, whereas the proportion of phospholipids is somewhat lower in the *glp-1(-)* mutant. When considering major lipid groups, the long-lived *glp-1(-)* and *fem-3(-)* mutants show higher Phosphatidic Acid (PA), but lower Sphingosine (So), compared to the *mog-3(-)* mutant. *glp-1(-)* mutant lacks germ cells, *fem-3(-)* mutant has occytes, and *mog-3(-)* mutant has hundreds of sperms, which may account for some of the differences in lipids, especially those that are membrane constituents of germ cells. Whether any of the lipid molecules that are differentially expressed in the long-lived and short-lived sterile mutants play a role in lifespan determination will require further investigation.

Our transcriptomic analysis is unique as we focused on probing the gene expression changes only in the intestine, which has been shown to be a key tissue to respond to germline signaling to mediate longevity. Since we dissected intestines from the different strains, we also avoided the caveat of comparing gene expression among strains with substantially different cell make-up. Overall, the transcriptomic data are consistent with the phenotypic data, as lacking sperms (*fem-3* mutant) or oocytes (*mog-3* mutant) results in gene expression changes that are part of those associated with a complete loss of germline (*glp-1* mutant) (Fig. 3 and Supplementary Fig. 3-5). This finding is entirely consistent with the notion that a complete lack of germline likely represents a much greater disruption to the organism, which triggers substantially greater gene expression changes ^13,31^.

The most prominent changes revealed by the RNA-seq analysis include fat metabolism and stress response genes. Expression change in fat metabolism genes is to be expected as the intestine represents the major metabolic tissue of *C. elegans* and our lipidomic analysis revealed greatly elevated TAGs and phospholipids in the mutants. Interestingly, among the lipid metabolism genes that showed significant changes, the changes detected in *fem-3(-)* and *mog-3(-)* each represent a subset of those detected in *glp-1(-)*. As discussed above, the absence of the entire germline likely elicits a greater need for metabolic reprogramming compared to lacking either sperms or oocytes, hence the substantially greater transcriptional changes detected in the *glp-1(-)* worms.

The *glp-1(-)* mutant is well-established to exhibit substantial gene expression changes in stress response genes. We now revealed that stress response genes are also over-represented among genes that show significant expression change in the feminized and masculinized mutants. This finding indicates that self-sterility, regardless of the precise cause, results in a heightened stress response. Interestingly, all three self-sterile mutants exhibit increased resistance to PA14 infection (Supplementary Fig. 6b). Therefore, even though “pathogen” genes are not over-represented among the differentially expressed genes in the *fem-3(-)* mutant, it nevertheless exhibits resistance to pathogen infection, similar to the *glp-1(-)* and *mog-3(-)* mutants.

Besides metabolism and stress, ribosome subunits are greatly over-represented among the differential expressed genes in *glp-1(-)* and *mog-3(-)*. At this time, the biological significance of these gene changes is not clear, but they appear to correlate with a lack of oogenesis. The *glp-1(-)* mutant also uniquely shows altered expression of cuticle/collagen genes. While collagen genes are generally thought to maintain the cuticle integrity, specific collagen genes have been demonstrated to have a major effect on longevity ^21,32^. Therefore, it is likely that altered expression of the many collagen genes in the *glp-1(-)* mutant contributes to its robust longevity extension.

Integrated analysis of our lipidomic and transcriptomic data highlighted a coordinated alteration of the sphingolipid metabolism pathway. Although all three sterile worms showed significant changes in the sphingolipid network, the exact genes and lipid molecules and their degree of change were somewhat different among the mutants. Interestingly, some of the changes, such as sphingosine level, appear to correlate with the longer-lived *glp-1(-)* and *fem-3(-)* as compared to the shorter-lived *mog-3(-)*. Future investigation of whether specific alterations of the sphingolipid pathway could impact longevity may provide further insights.

Despite that all three sterile mutants upregulate many stress response genes, they exhibit very different lifespans under normal growth conditions. Since DAF-16 and SKN-1 are two of the best-characterized transcription factors that play a key role in longevity, stress response, and fat metabolism in *C. elegans* ^6,21,28,33^, we further investigated their activities in the sterile mutants. Our results supported that SKN-1 is activated in all three sterile mutants, as indicated by the induction of GST-4, a well-established downstream target of SKN-1, and the requirement of SKN-1 for the full longevity of the sterile mutants. Furthermore, our results suggested that SKN-1 activation requires the accumulated fat, consistent with earlier studies indicating that excess fat and lipid toxicity activate SKN-1^21^. Interestingly, while SKN-1 activation contributed to the full lifespan of the sterile mutants, as well as the wild-type control, the degree and kinetics of its activation (Fig 5 and 6, Supplementary Fig. 7-10) did not appear to correlate with the overall longevity.

On the contrary, we observed that DAF-16 is activated in the longer-lived mutants *glp-1(-)* and *fem-3(-)*, as indicated by DAF-16 nuclear localization in the intestine, but not in the shorter-lived *mog-3(-)* mutant. DAF-16 is one of the best-characterized longevity-promoting factors in *C. elegans*, and its activation in the long-lived sterile strains is fitting. We were curious to know what signals for DAF-16 activation. Using *fat-6* RNAi to deplete lipid storage, we found that DAF-16 activation was partially attenuated, suggesting that DAF-16 activation in *glp-1(-)* and *fem-3(-)* partially depends on the accumulated fat. However, since DAF-16 is not activated in *mog-3(-)* mutant despite its excess lipid accumulation, other factors must contribute to DAF-16 activation. It remains possible that the complete absence of sperm differentiation in the *glp-1(-)* and *fem-3(-)* mutants induce a signal to activate DAF-16 in the intestine, a hypothesis that requires further testing. Interestingly, whereas DAF-16 is similarly activated in both *glp-1(-)* and *fem-3(-)* mutants, *glp-1(-)* has a greater lifespan extension compared to *fem-3(-)*. Therefore, additional factors likely contribute to the greater longevity of the *glp-1(-)* mutant.

## Methods

### Oil red O (ORO) staining

The ORO staining procedure was carried out as previously described with slight modifications^34^. A 5 mg ORO stock solution was prepared by dissolving the ORO (Sigma-Aldrich, O0625) in 1 ml of isopropanol and left to equilibrate for 2 days on a magnetic stirrer. ∼100 worms were collected in 500 μl PBST (PBS + 0.1% Tween 20), washed once with PBS, and fixed in 4% formaldehyde fixative (800 μl water, 100 μl 10X PBS, and 100 μl formaldehyde) for 30 minutes. After removal of the fixative, the worms were washed with PBST and PBS once each, and then treated with 60% isopropanol for 2 minutes. 200 μl working solution of ORO (60% ORO stock + 40% water) was added to each tube and incubated for 2-3 hours. The worms were washed twice with PBS, and 10 μl of Vectashield mounting media was added to each tube. For imaging, the worms were mounted on a 2% agarose pad and images were captured using a Leica compound microscope (Leica DM5000B).

### ORO extraction and calorimetric estimation

All steps of the ORO staining protocol were meticulously followed up until the first wash with PBS. The worms were washed once with 500 μl of PBS and twice with distilled water for 5 minutes each. Subsequently, they were incubated twice in 60% isopropanol for 5 minutes each. To extract the bound ORO, the worms were placed in 100 μl of 100% isopropanol and incubated for 5-7 minutes on a rocking platform. 85-90 μl of the supernatant was then transferred to a 96-well plate and the absorbance was measured at 492 nm. A 100% isopropanol control was utilized to subtract the background signal.

### RNAi treatment

The RNAi procedure was carried out as previously described with slight modifications^35^. The RNAi was conducted by feeding the worms with HT115 bacteria that expressed double-stranded RNA of the target gene. The RNAi clones were obtained from RNAi feeding libraries created by Julie Ahringer and Marc Vidal’s laboratories^36,37^.

To begin, a single colony of RNAi bacteria was streaked onto an LB agar plate containing ampicillin (100 μg/mL) and grown overnight at 37°C. The following day, a single bacterial colony was inoculated in 5 mL of LB broth containing ampicillin (100 μg/mL) and tetracycline (15 μg/mL) and incubated overnight in a shaker at 37°C. To verify the accuracy of the RNAi clone, the plasmid was extracted and submitted for sequencing. Upon confirmation of the correct RNAi plasmid insert, a single colony of the bacteria was cultured overnight in 5 mL of LB broth containing ampicillin (100 μg/mL) and tetracycline (12.5 μg/mL) at 37°C in a shaker. For every RNAi experiment, a control culture of bacteria containing the L4440 plasmid was also included.

The overnight culture (5% of the final culture) was then inoculated in the desired volume of LB broth containing ampicillin (100 μg/mL) and incubated for an additional 3-4 hours at 37°C in a shaker. Once the optical density (OD^600^) reached approximately 0.6, the culture was induced with 1 mM IPTG (Isopropyl β-D-1-thiogalactopyranoside) and incubated for an additional 2-3 hours at 37°C in a shaker.

The culture was then centrifuged at 3000 RPM for 15 minutes at room temperature, and the bacterial pellet was resuspended in the desired volume of the same supernatant (200 μl per plate). Two or three RNAi plates were prepared for each target gene in each experiment, each containing 200 μL of bacterial culture spread on NGM agar plate (no streptomycin) containing ampicillin (100 μg/mL), tetracycline (15 μg/mL), and IPTG (1 mM). The plates were dried under a hood, and 10-15 adult worms were allowed to lay eggs overnight on each plate. The worms from the first set of RNAi plates were transferred to a new set of RNAi plates, and the embryos were collected for 2-3 hours. The embryos laid on the second set of RNAi plates were grown to the desired stage, and then the relevant phenotypes were scored.

### Lifespan Analysis

Lifespan assays were performed with some modifications as described earlier^35^. 10-15 gravid worms were placed on plates containing the desired bacterial food, and approximately 200 embryos were collected. Upon reaching the 1-day adult stage, 30 worms were transferred to each of three RNAi plates and their status was monitored every other day by counting the number of live or dead worms. Death was confirmed by the absence of movement after gentle touching of the worm’s nose. Cases of death by rupturing, bursting, or runoff were considered censored. The data was analyzed using the OASIS online survival analysis tool^38^ available at https://sbi.postech.ac.kr/oasis2/.

### TAG (Triacyglycerides) estimation

Approximately 100 worms were collected in PBST, washed three times, and then suspended in 100 μL of cold PBST. The mixture was flash-frozen in liquid nitrogen and thawed on ice. The worms were homogenized using a motorized pestle (Kontes: 749540-0000) and centrifuged for 3 minutes at 13,000 g at 4°C. A portion (10-20 μL) of the supernatant was set aside for protein measurement and were frozen and stored at -80°C. The remaining supernatant was heated at 70°C for 10 minutes. 10-25 μL of the heated sample (in two replicates) and 200 μL of infinity triglyceride reagent (ThermoFisher, TR22421) were added to separate wells of a 96-well plate and incubated at 37°C for 30 minutes. The plate was wrapped in aluminum foil for the duration of the experiment, and absorbance was measured at 540 nm.

### *Pseudomonas aeruginosa* (PA14) slow-killing and survival assays

The PA14 slow-killing assay was modified from the method described earlier^24^. A single colony of PA14 was inoculated into LB media and cultured overnight at 37°C in a shaker. The following day, the culture was concentrated 20-fold and spread on PA14 slow-killing plates, then dried under a hood.

Approximately 10-15 gravid worms were placed on the PA14 plate, and around 200 embryos were collected. When the worms reached the L4 stage, 50 μM FUDR (5-fluoro-2′-deoxyuridine) was added to both the control and experimental PA14 plates to inhibit reproduction. The following day, 30 worms were transferred to each of three PA14 plates and the number of live or dead worms were scored every 12 hours until all worms had died. Death by rupture, bursting, or crawling off the plates was considered censored. The data was analyzed using the OASIS (https://sbi.postech.ac.kr/oasis2/) online survival analysis tool^38^.

### RNA isolation and library preparation for RNA-seq

Lifespan assays were performed as previously described with some modifications^35^. For the collection of whole worm samples, approximately 200 worms were collected in TRI reagent. For the collection of intestinal tissue, 100-120 worms were dissected using a syringe needle, and 70-80 intestines were collected in TRI reagent. The samples were homogenized in 10 volumes of TRI reagent (500 μL) by performing several rounds of freeze-thaw cycles in liquid nitrogen. Subsequently, 200 μL of chloroform was added per 1 mL of TRI reagent (MRC, TR118), the tube was vortexed for 15 seconds, and the mixture was incubated at room temperature for 2-3 minutes. The samples were then centrifuged at 12000 g for 15 minutes at 4°C in a refrigerated centrifuge. The upper colorless layer was collected in a 1.5 mL microcentrifuge tube and 0.5 mL of isopropanol per mL of TRIZOL was added. The tube was mixed well and the sample was incubated at room temperature for 10 minutes. The tube was then centrifuged at 12000 g for 10 minutes at 4°C. The supernatant was removed, and the transparent RNA pellet was washed with 1 mL of 75% ethanol. After removing all residual ethanol, the pellet was air-dried for 5-7 minutes at room temperature and the RNA was dissolved in RNase-free water. The RNA was then purified using the RNeasy Mini Kit (Qiagen, 74106). A library for RNA-seq was prepared using the Ovation Human FFPE RNA-Seq Library System (NuGEN), and *C. elegans*-specific oligos were used to remove small RNAs.

### RNA-seq data analysis

Transcriptomic data analysis was performed using the pipeline as described earlier^35,39^. Low quality reads were filtered out using FASTX toolkit and adaptors sequences were trimmed using cutadapt. tRNA and rRNA were removed using bowtie2 and remaining reads were mapped to WS250 (wormbase release 250) using Tophat2. Raw gene counts were calculated using Cufflinks and differential expression of each transcript was analyzed using EdgeR under R studio. The average transcript counts of two replicates under 10 were filtered out and RPKM was analyzed for the filtered set of genes. A 5% false discovery rate (FDR) was used to determine significant differential expression. Heatmaps were generated using Morpheus web-based tool. (https://software.broadinstitute.org/morpheus/) and Edge R. Principal component analysis (PCA) was performed using EdgeR^15^.

### Sample preparation for LC-MS-based Lipidomic analysis

20-25 gravid worms were placed on ten 6-cm NGM plates that were seeded with OP50 bacterial food. Embryos were then collected and when they reached the 1-day adult stage, 1,500 to 2,000 worms were picked and placed on a seeded plate. The worms were then washed off the plate using M9 buffer + 0.05% Tween-20 in a 5 ml Falcon tube. They were further washed once with M9 buffer and double-distilled water. The worms were quenched with 500 μl of cold (−20°C) MeOH and the sample was flash-frozen in liquid nitrogen and stored at -80°C before further processing. The sample was then thawed on ice, 1.7 ml of Methyl tert-butyl ether (MTBE) was added, and the sample was vigorously vortexed. The sample was then handed over to the Cornell Metabolomics facility for further processing and Liquid Chromatography-Mass Spectrometry (LC-MS) analysis.

### Lipidomic Data Analysis

Differential expression of each lipid molecule ‘features’ was analyzed using EdgeR under R studio. A 5% false discovery rate (FDR) was used to determine significant differential changes. Heatmaps were generated using Morpheus web-based tool. (https://software.broadinstitute.org/morpheus/). Principal component analysis (PCA) was performed using EdgeR^15^.

### Lipidomic data set enrichment analysis

Lipidomic data set enrichment analysis was analyzed using MetaboAnalyst (www.metaboanalyst.ca). It is a web-based server designed for metabolomic data analysis and omic data integration. The differentially produced lipid molecules were uploaded on the pathway analysis module and screened for enriched lipid pathways.

### Gene ontology analysis

Gene ontology (GO) term analysis was performed using the WormCat ^22^.

### STRING analysis

STRING analysis was performed using an online tool, STRING, which is designed for Protein-Protein Interaction and Functional Enrichment Analysis^23^. A list of differentially regulated genes was uploaded to the platform and was clustered using k-means clustering based on their Gene Ontology terms.

### Omics data integration

The integration of transcriptomic and lipidomic data was performed using MetaboAnalyst (www.metaboanalyst.ca). This module was introduced in MetaboAnalyst 3.0. In this module, the lists of differentially expressed genes and differentially produced lipids were uploaded with their log2 fold change. The degree of integration between these two data sets was analyzed. Furthermore, genes and lipids were mapped to relevant KEGG metabolic pathways for over-representation analysis.

## Acknowledgements

We thank C. elegans Genetic Center (CGC) for providing worm strains which is funded by the NIH Office of Research Infrastructure Programs (P40 OD010440). We thank Frank C. Schroeder laboratory (BTI institute, Cornell) especially Frank C. Schroeder, Pooja Gudibanda and Bennett William Fox for the insightful discussions and suggestions. We thank Lee laboratory members for meaningful suggestions.

We thank Felicity J. Emerson (Lee Lab) for reading the manuscript. We thank Amy J. Walker (UMass Chan Medical School) for the initial help in Gene Ontology term analysis. We thank the Genomics Facility (RRID:SCR_021727) and Proteomics and Metabolomics facility (RRID:SCR_021743) of the Biotechnology Resource Center (BRC) of Cornell Institute of Biotechnology for RNA-sequencing and lipidomic experiments, respectively. This work was supported by NIA grant AG024425 to SSL.

## Author’s contributions

AC and SSL conceptualized the study and designed the experiment. AC performed the experiments. AC and SSL interpreted the data and wrote the manuscript. SSL supervised the study and acquired the funding.

