## Supplementary Figures for "Different gametogenesis states uniquely impact longevity in *Caenorhabditis elegans*"

**Supplementary Information for Chaturbedi and Lee**

Pages

Supplementary Table Details	2

Supplementary Figures	3-13

**Supplementary Table Details:**

| Supplementary Table 1 | Lifespan data and statistics shown in Supplementary Fig. 1. |
| --- | --- |
| Supplementary Table 2 | Lifespan data and statistics shown in Fig. 1. |
| Supplementary Table 3 | List of lipid molecules detected in lipidomic analysis and list of significantly differentially-produced lipid molecule in different genotype pairs. |
| Supplementary Table 4 | List of overlapping upregulated genes between *glp-1(-)*_intestine, *glp-1(-)*_whole-worm and *glp-1(-)*_Steinbaugh et al., 2015 (relative to wild-type). |
| Supplementary Table 5 | List of significantly up- and down-regulated genes in different genotype pairs. |
| Supplementary Table 6 | List of overlapping upregulated genes between *fem-3(-) (*relative to wild-type) in whole-worm and intestinal data-sets. |
| Supplementary Table 7 | List of overlapping upregulated genes between *mog-3(-) (*relative to wild-type) in whole-worm and intestinal data-sets. |
| Supplementary Table 8 | List of overlapping upregulated genes between *glp-1(-)*, *fem-3(-)* and *mog-3(-)* worms (relative to wild-type). |
| Supplementary Table 9 | List of enriched Gene Ontology terms in the upregulated gene-set in *glp-1(-)*, *fem-3(-)* and *mog-3(-)* worms (relative to wild-type). |
| Supplementary Table 10 | List of significantly up- and down-regulated fat metabolism genes in different genotype pairs. |
| Supplementary Table 11 | List of significantly up- and down-regulated stress response genes in different genotype pairs. |
| Supplementary Table 12 | List of enriched Gene Ontology terms and genes in each cluster in the upregulated gene-set in *glp-1(-)*, *fem-3(-)* and *mog-3(-)* worms (relative to wild-type) using STRING. |
| Supplementary Table 13 | List of enriched Gene Ontology terms and genes in each cluster in the upregulated fat metabolism genes in *glp-1(-)*, *fem-3(-)* and *mog-3(-)* mutant worms (relative to wild-type) using STRING. |
| Supplementary Table 14 | List of overlapping upregulated pathogen response genes between *glp-1(-)*, *fem-3(-)* and *mog-3(-)* mutant worms (relative to wild-type). |
| Supplementary Table 15 | Survival data and statistics of *glp-1(-)*, *fem-3(-)* and *mog-3(-)* worms upon *Pseudomonas aeruginosa* (PA14) infection. |
| Supplementary Table 16 | Lifespan data and statistics shown in Fig. 5. |

**Supplementary Fig. 1**

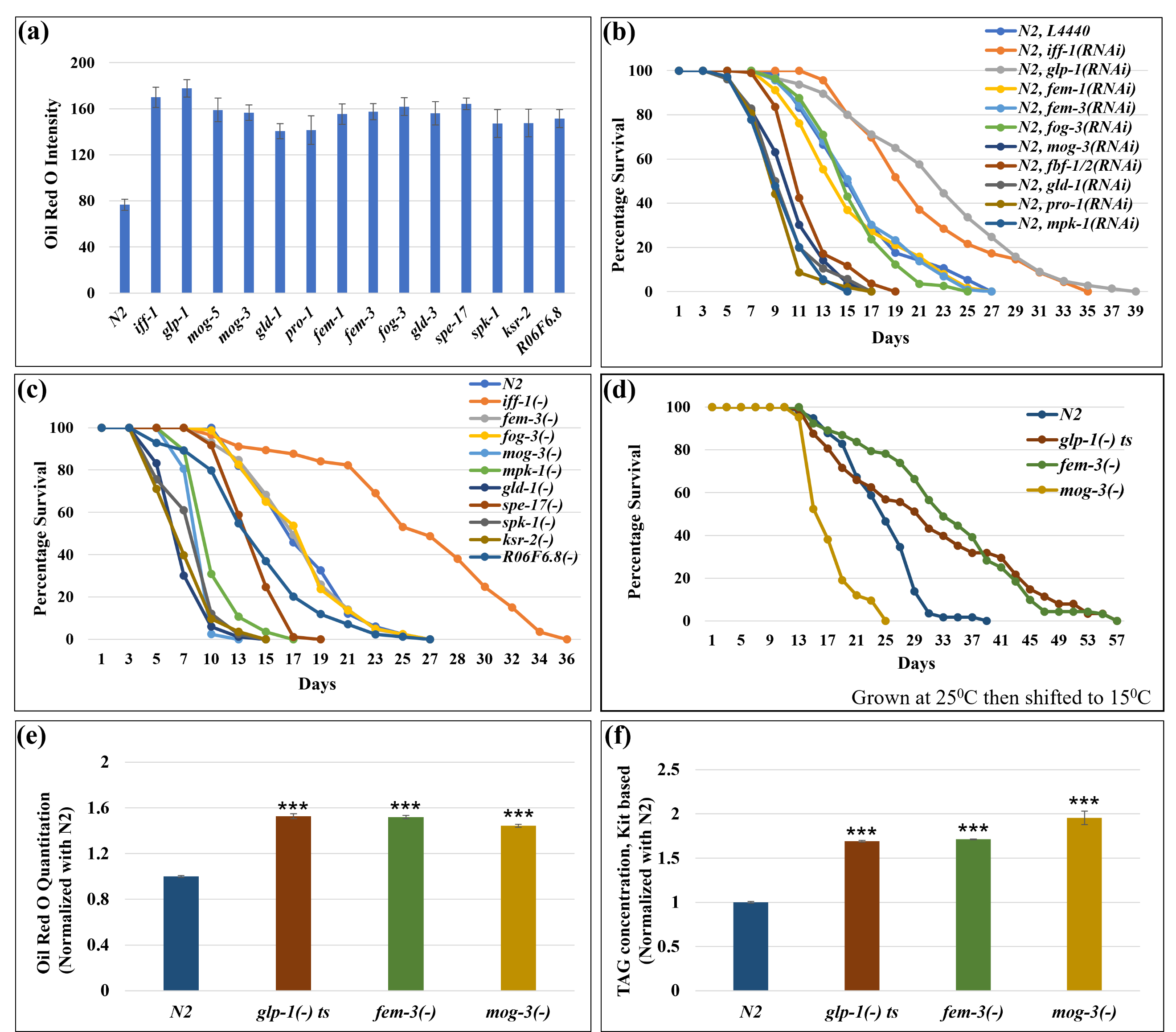

**Supplementary Fig. 1: Sterile strains displayed excessive lipid accumulation but different lifespans.**

(a). Oil red O (ORO) intensity in the indicated sterile strains (day 1 adults). All three sterile mutants accumulated excess fat. ORO intensity is shown as the mean intensity of two replicates for each genotype. n>10 for each replicate.

(b, c, d). Survival curves showing the lifespans (at 20^o^C) of wild-type worms treated with the indicated gene RNAi (b), the lifespans (at 20^o^C) of wild-type and mutant strains (c), the lifespans of wild-type N2, *glp-1(e2141)*, *fem-3(e1996)* and *mog-3(q74)* worms grown at 25^o^C then aged at 15^o^C (d). Lifespan data and statistics are shown in Supplementary Table 1.

Sterile *glp-1(e2141)* worms lived longer, and *mog-3(q74)* worms lived shorter irrespective of the temperature at they were grown during adulthood. *fem-3(e1996)* worms lived longer when grown at 25^o^C.

For lifespan assays, at least 90 worms were scored for each genotype per replicate and two independent experiments were performed for each genotype.

(e, f). Lipid levels in wild-type N2, *glp-1(e2141)*, *fem-3(e1996)* and *mog-3(q74)* worms (day 1 adults) based on the optical density of extracted Oil Red O from 100 worms (e), and calorimetry-based triacylglycerides (TAG) kit (normalized with N2) (f). ORO and TAG levels were quantified from at least 100 worms for each replicate and two replicates were performed for each assay. ***p < 0.001.

**Supplementary Fig. 2**

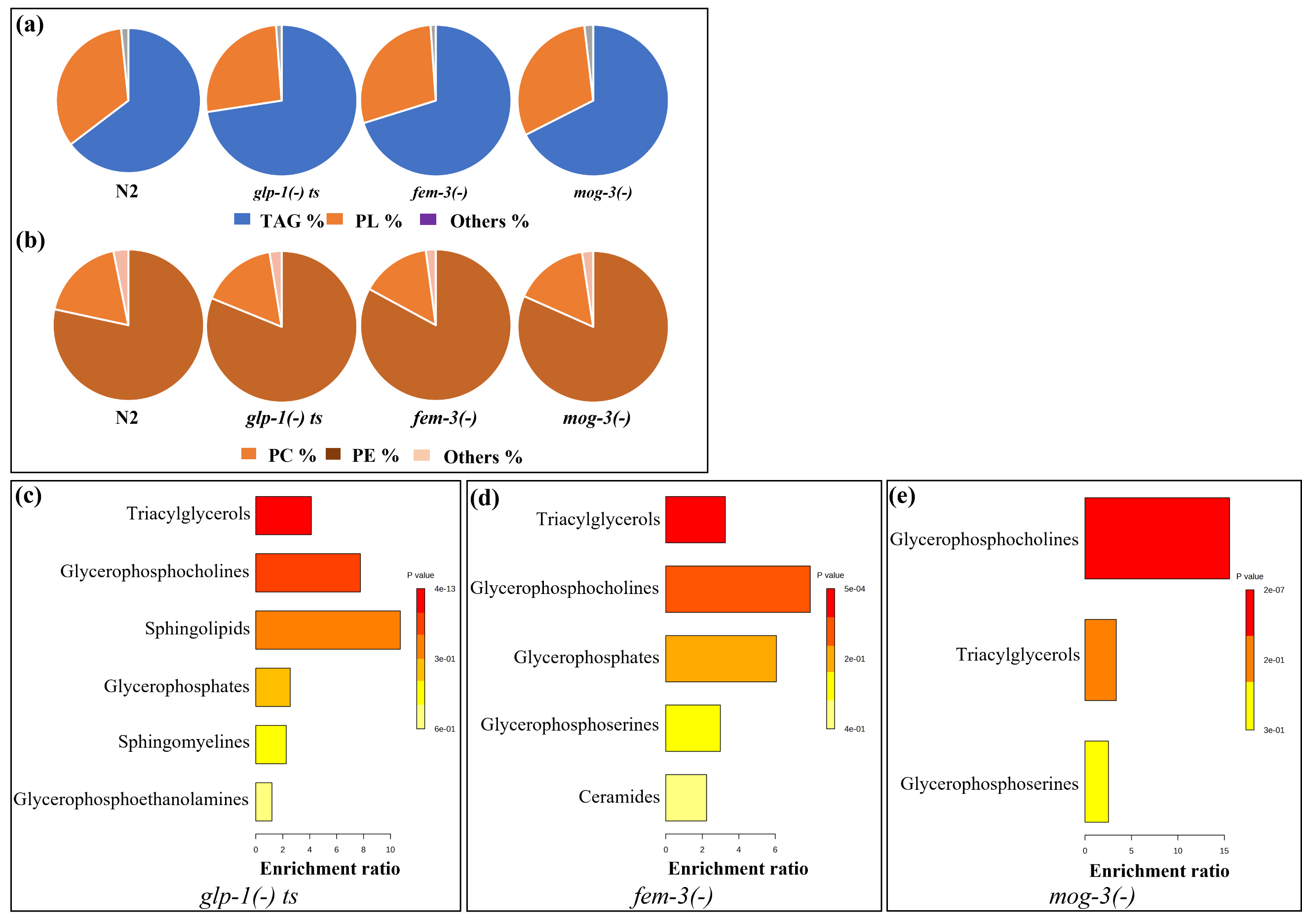

**Supplementary Fig. 2: Different sterile mutants displayed similar lipid profiles.**

a) Relative distribution of total lipids in wild-type N2, *glp-1(e2141)*, *fem-3(e1996)* and *mog-3(q74)* worms. The large majority of *C. elegans* lipidome was comprised of Triacylglycerides (TAG) and Phospholipids (PL). All three sterile mutants stored increased levels of TAG compared to wild-type.

(b). Relative distribution of phospholipids in wild-type N2, *glp-1(e2141)*, *fem-3(e1996)* and *mog-3(q74)* worms. Phospholipids were mainly stored in the form of Phosphatidylcholine (PC) and Phosphatidylethanolamine (PE) and their relative distribution stayed similar in the indicated genotypes.

(c, d, e). The lipid molecules that were upregulated in *glp-1(e2141)* (c)*, fem-3(e1996)* (d) and *mog-3(q74)* (e) (compared to wild-type) were subjected to lipid group enrichment analysis using MetaboAnalyst, a web-based platform for metabolomic/lipidomic data analysis that conducts enrichment analysis based on several libraries containing mammalian metabolite sets.

**Supplementary Fig. 3**

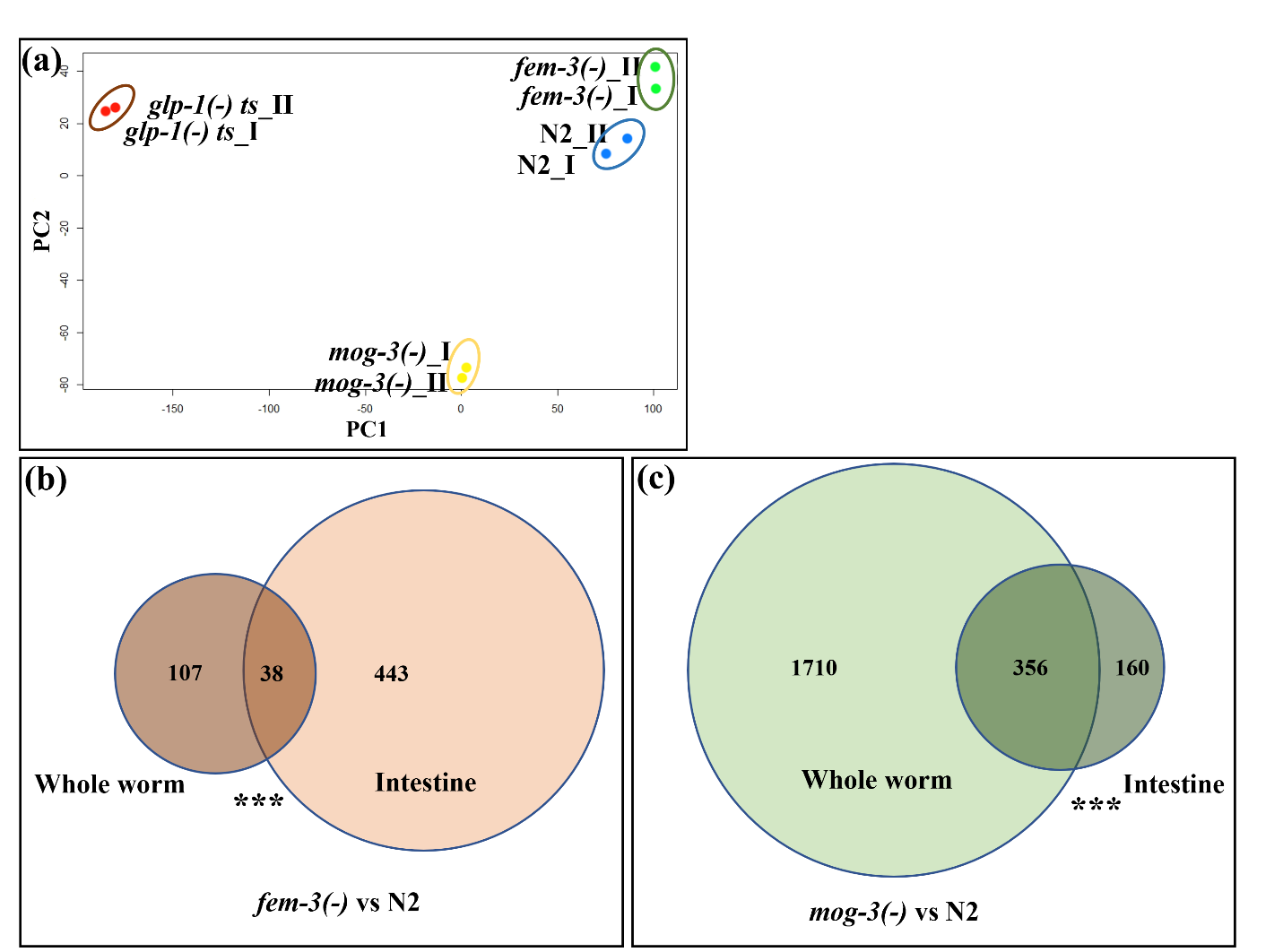

**Supplementary Fig. 3**: **Whole-worm transcriptomic analysis of the sterile mutants.**

(a). Principal components analysis (PCA) of the whole-worm RNA-seq data for two replicates of N2, *glp-1(e2141)*, *fem-3(e1996)* and *mog-3(q74)* worms. Each dot indicates one replicate. Long-lived *glp-1(e2141)* and *fem-3(e1996)* worms were clearly separated from short-lived *mog-3(q74)* worms on PC2. PCA plot was generated using EdgeR.

(b, c). Venn diagrams showed the significant overlaps between the upregulated genes in *fem-3(e1996)* (b) (detailed list of genes shown in Supplementary Table 6) and *mog-3(q74)* (c) (detailed list of genes shown in Supplementary Table 7) relative to N2, in whole-worm and intestinal data sets. Venn diagrams were generated using EdgeR. ***p < 0.001.

**Supplementary Fig. 4**

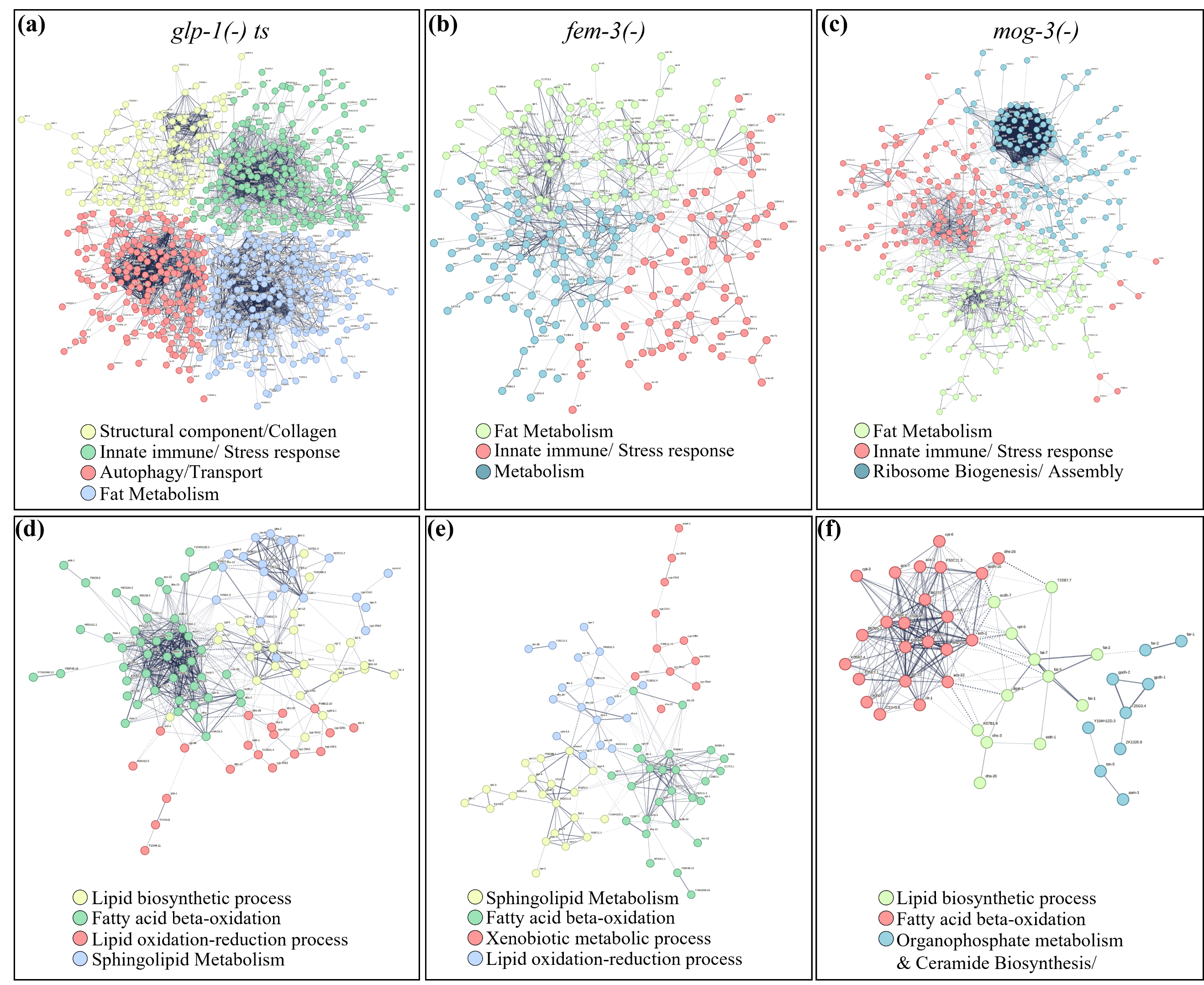

**Supplementary Fig. 4:** **STRING analysis of the genes that were differentially expressed in the sterile mutants.**

STRING is a web-based tool that clusters proteins based on a variety of data, including protein-protein interaction, subcellular localization, and more^23^ (a, b, c). K-means clustering showing the major groups of upregulated genes in *glp-1(e2141)* (a)*, fem-3(e1996)* (b) and *mog-3(q74)* (c) vs N2. The detailed list of genes in each cluster is shown in Supplementary Table 12).

(d, e, f). K-means clustering showing the subgrouping of the upregulated fat metabolism genes (as derived from a-c) in *glp-1(e2141)* (d)*, fem-3(e1996)* (e) and *mog-3(q74)* (f) vs N2. The detailed list of genes in each cluster is shown in Supplementary Table 13).

**Supplementary Fig. 5**

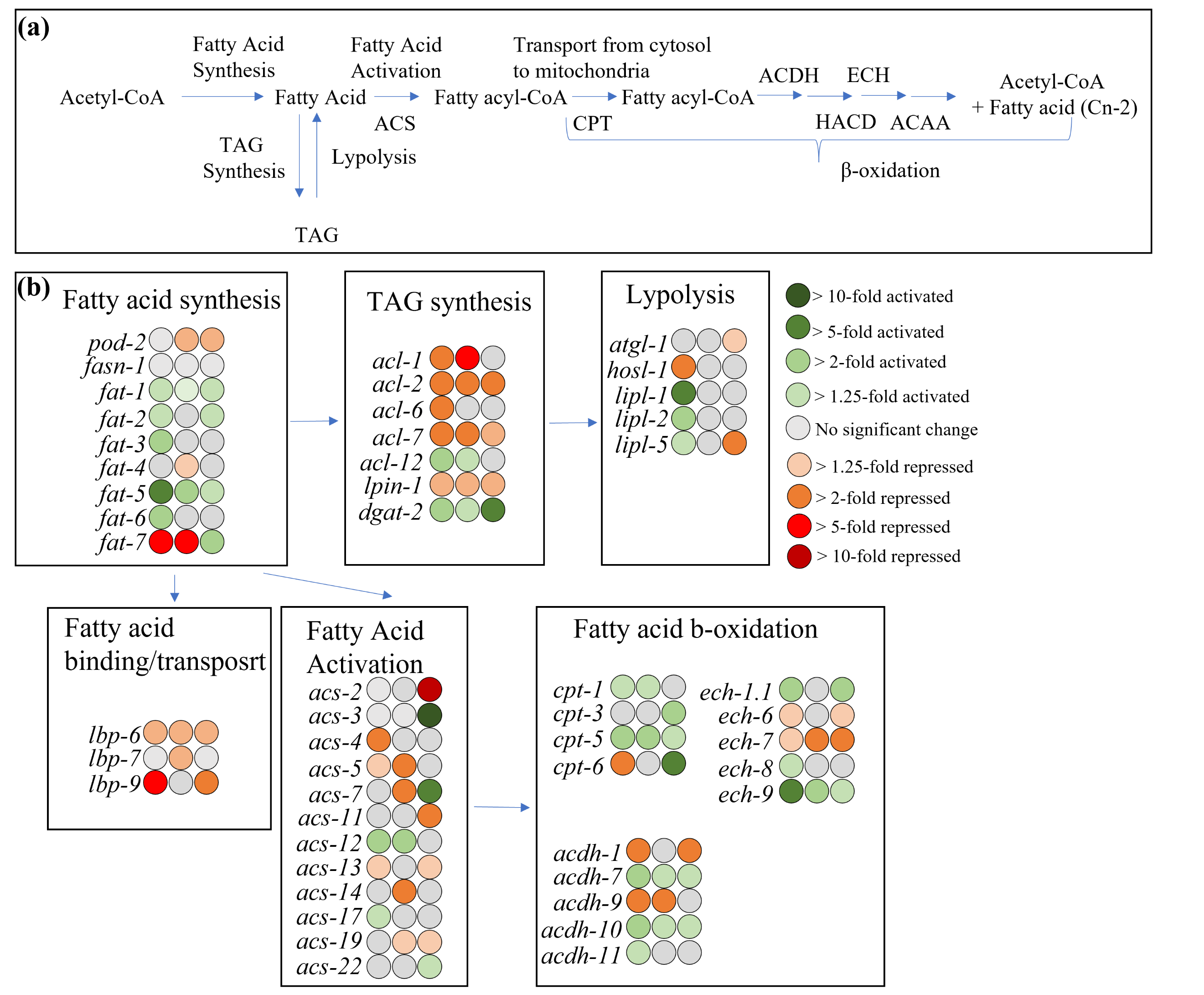

**Supplementary Fig. 5:** **Fold change of specific fat metabolism genes in the sterile mutants.**

(a). Schematic diagram showing the major enzymes in the fatty acid synthesis and degradation pathway. (b) Fold change in RNA expression for the indicated genes in *glp-1(e2141)* (left)*, fem-3(e1996)* (middle) and *mog-3(q74)* (right) vs N2 are represented according to the color scale.

**Supplementary Fig. 6**

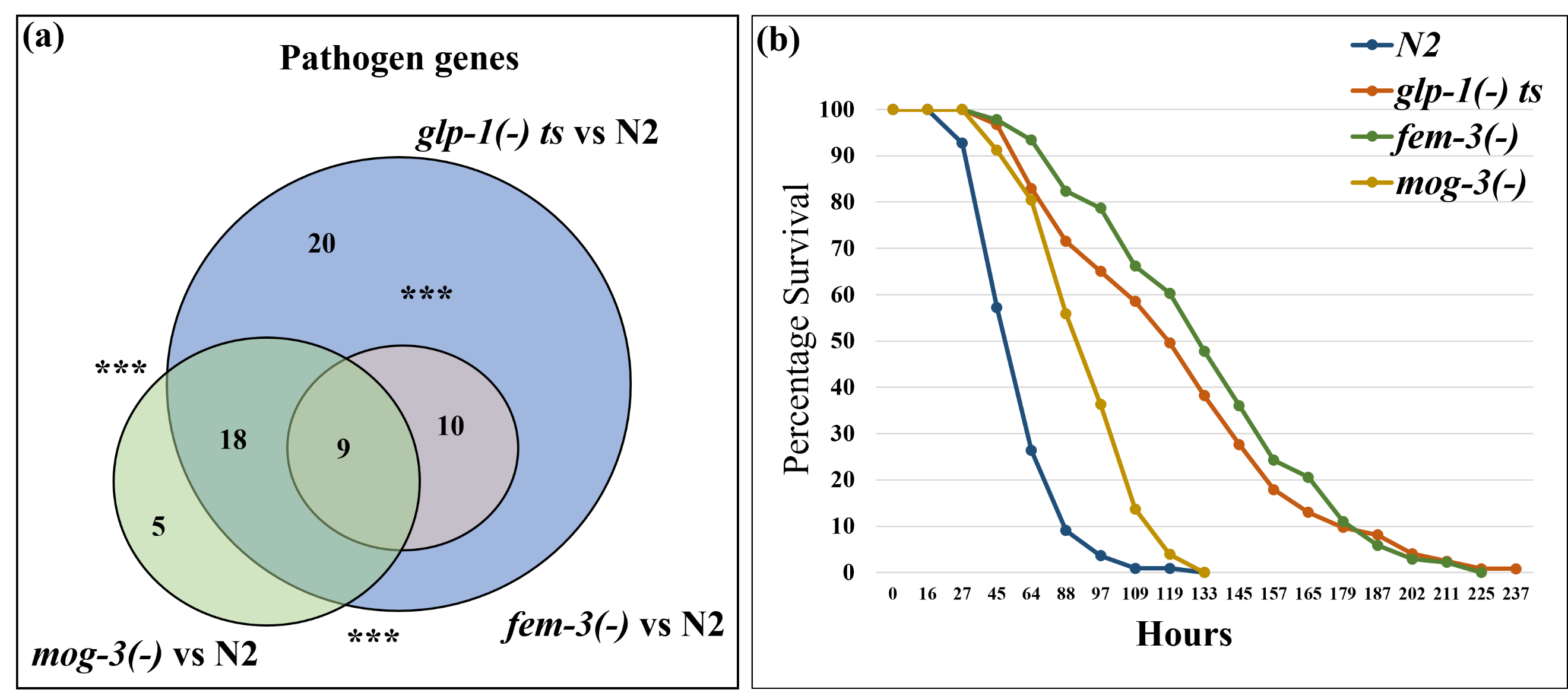

**Supplementary Fig. 6:** **Sterile mutants are less susceptible to *Pseudomonas aeruginosa* (PA14) infection.**

(a). Venn diagram showing the overlaps between the upregulated pathogen response genes in *glp-1(e2141), fem-3(e1996),* and *mog-3(q74)*. The Detailed lists of genes and overlaps are shown in Supplementary Table 14. Venn diagrams were generated using edgeR (Robinson et al., 2010). ***p < 0.001.

(b). Survival curves showing the lifespans of N2, *glp-1(e2141)*, *fem-3(e1996)* and *mog-3(q74)* worms upon PA14 infection in a slow-killing assay at 25^o^C. Worms were exposed to PA14 starting at day 4 of adulthood to avoid death of wild-type N2 worms from “bagging” (progeny hatching inside the uterus). All three sterile strains showed better survival upon PA14 infection (survival data and statistics shown in Supplementary Table 6). For lifespan, at least 90 worms were scored for each genotype per replicate and two independent experiments were performed for each genotype.

**Supplementary Fig. 7**

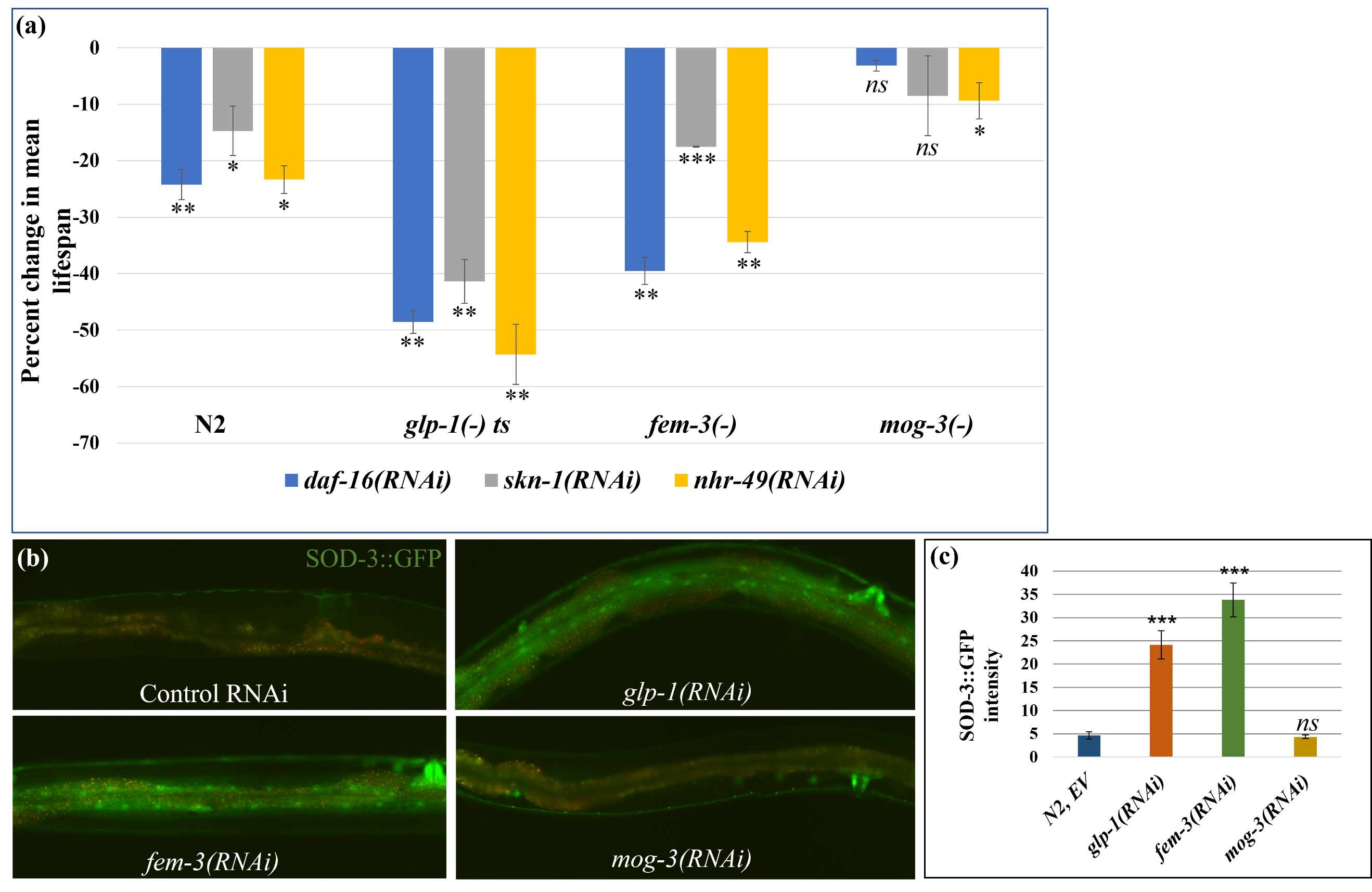

**Supplementary Fig. 7:** **The lifespans of different sterile mutants are dependent on distinct genetic pathways.**

(a). Percent change in mean lifespans of N2, *glp-1(e2141)*, *fem-3(e1996)* and *mog-3(q74)* worms upon RNAi depletion of *daf-16, skn-1* and *nhr-49* (survival curves shown in Figure 4, a-d). Percent change in mean lifespan shown here represents the average of two replicates. At least 90 worms were scored for each genotype for every replicate. * p <0.05, ** p<0.01, ***< 0.001, ns – not significant.

(b, c). Representative images (d) and quantification (e) showed that nuclear localization of SOD-3::GFP is elevated in *glp-1(RNAi)* and *fem-3(RNAi)* worms but not in *mog-3(RNAi)* worms. Y-axis represents mean GFP intensity of two replicates of each genotype with at least 10 worms per replicate. ***p < 0.001, ns - not significant.

**Supplementary Fig. 8**

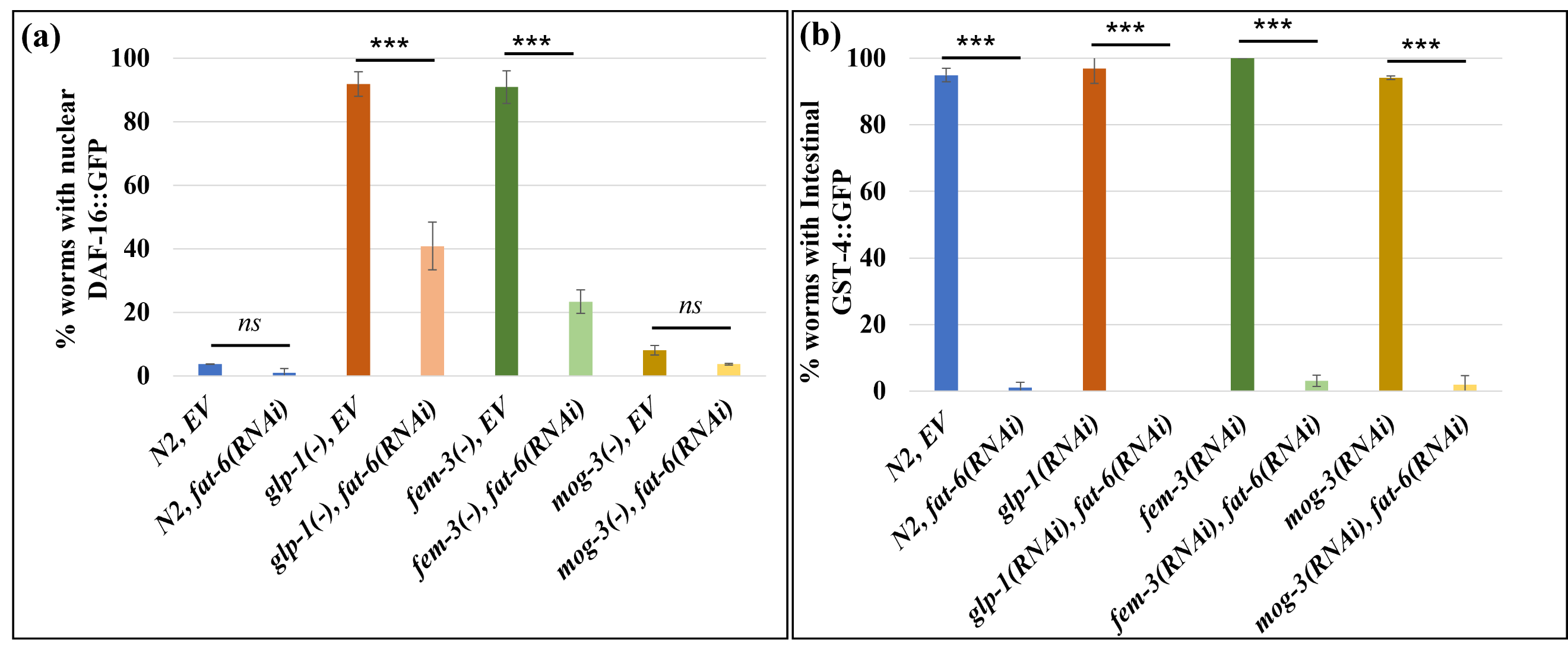

**Supplementary Fig. 8: DAF-16::GFP and GST-4::GFP activation are differentially dependent on available fat.**

a) Percent worms with nuclear DAF-16::GFP in the indicated genotypes. RNAi depletion of *fat-6* partially attenuated DAF-16::GFP nuclear localization. b) Percent worms with intestinal GST-4::GFP in the indicated genotypes. RNAi depletion of *fat-6* completely abolished intestinal GST-4::GFP expression.

The percent shown here is the mean of two replicates (at least 50 worms per replicate) of the indicated genotypes. These experiments were performed at 25^o^C.

***p < 0.001, ns - not significant.

**Supplementary Fig. 9**

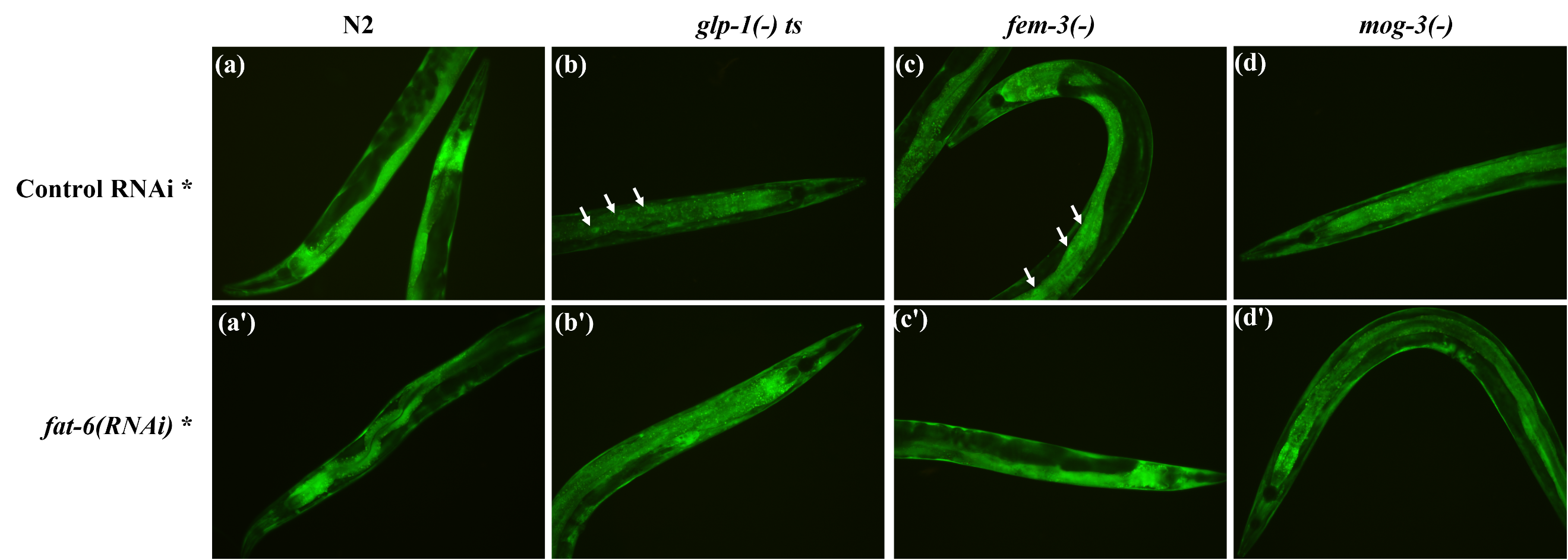

**Supplementary Fig. 9: DAF-16 nuclear localization is partially dependent on available fat.**

(a, b, c, d). The expression of DAF-16::GFP in N2 (a), *glp-1(e2141)* (b)*, fem-3(e1996)* (c) and *mog-3(q74)* (d) worms. Arrowheads show nuclear GFP in *glp-1(e2141)* and *fem-3(e1996)* worms.

(a**'**, b**'**, c**'**, d**'**). The expression of DAF-16::GFP in in N2 (a**'**), *glp-1(e2141)* (b**'**)*, fem-3(e1996)* (c**'**) and *mog-3(q74)* (d**'**) worms upon RNAi depletion of *fat-6.* RNAi depletion of *fat-6* partially rescued nuclear GFP phenotype (only worms with non-nuclear GFPs are shown here). These experiments were performed at 25^o^C.

**Supplementary Fig. 10**

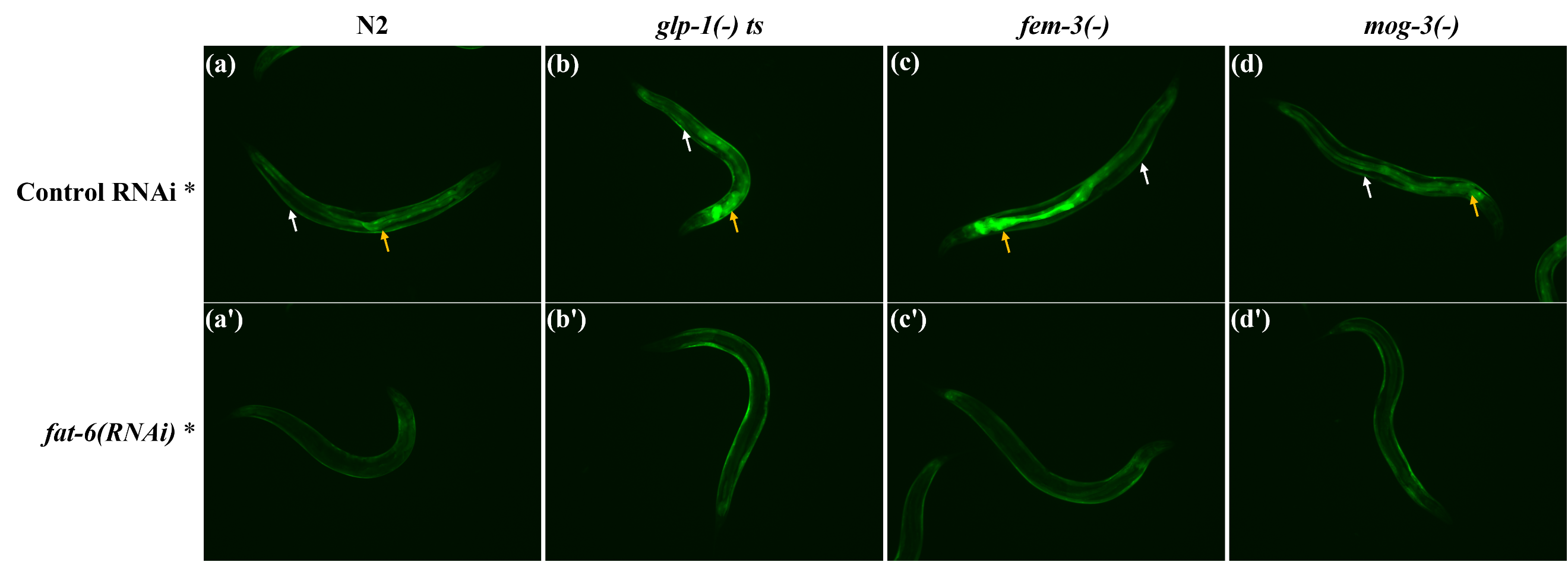

**Supplementary Fig. 10: Intestinal SKN-1 activation is completely dependent on available fat.**

(a, b, c, d). The expression of GST-4::GFP in N2 (a), *glp-1(e2141)* (b)*, fem-3(e1996)* (c) and *mog-3(q74)* (d) worms. White and yellow arrows show hypodermal and intestinal GFP expression, respectively. GST-4::GFP is preferentially activated in the intestines of sterile worms.

(a**'**, b**'**, c**'**, d**'**). The expression of GST-4::GFP in N2 (a**'**), *glp-1(e2141)* (b**'**)*, fem-3(e1996)* (c**'**) and *mog-3(q74)* (d**'**) worms upon *fat-6* RNAi. RNAi depletion of *fat-6* specifically deactivated intestinal GST-4::GFP expression.

These experiments were performed at 25^o^C.
